# A systematic pipeline for classifying bacterial operons reveals the evolutionary landscape of biofilm machineries

**DOI:** 10.1101/769745

**Authors:** Cedoljub Bundalovic-Torma, Gregory B. Whitfield, Lindsey S. Marmont, P. Lynne Howell, John Parkinson

## Abstract

In bacterial functionally related genes comprising metabolic pathways and protein complexes are frequently encoded in operons and are widely conserved across phylogenetically diverse species. The evolution of these operon-encoded processes is affected by diverse mechanisms such gene duplication, loss, rearrangement, and horizontal transfer. These mechanisms can result in functional diversification of gene-families, increasing the potential evolution of novel biological pathways, and serves to adapt pre-existing pathways to the requirements of particular environments. Despite the fundamental importance that these mechanisms play in bacterial environmental adaptation, a systematic approach for studying the evolution of operon organization is lacking. Herein, we present a novel method to study the evolution of operons based on phylogenetic clustering of operon-encoded protein families and genomic-proximity network visualizations of operon architectures. We applied this approach to study the evolution of the synthase dependent exopolysaccharide (EPS) biosynthetic systems: cellulose, acetylated-cellulose, poly-β-1,6-*N*-acetyl-D-glucosamine (PNAG), Pel, and alginate. These polymers have important roles in biofilm formation, antibiotic tolerance, and as virulence factors in opportunistic pathogens. Our approach reveals the complex evolutionary landscape of EPS machineries, and enabled operons to be classified into evolutionarily distinct lineages. Cellulose operons show phyla-specific operon lineages resulting from gene loss, rearrangement, and the acquisition of accessory loci, and the occurrence of whole-operon duplications arising through horizonal gene transfer. Our evolutionary-based classification also distinguishes between the evolution of PNAG production between Gram-negative and Gram-positive bacteria on the basis of structural and functional evolution of the acetylation modification domains shared by PgaB and IcaB loci, respectively. We also predict several *pel*-like operon lineages in Gram-positive bacteria, and demonstrate in our companion paper (BIORXIV/2019/768473) that *Bacillus cereus* produces a Pel-dependent biofilm that is regulated by cyclic-3’,5’-dimeric guanosine monophosphate (c-di-GMP).

**AUTHOR SUMMARY:** In bacterial genomes biological processes are frequently encoded as operons of co-transcribed neighbouring genes belonging to diverse protein families. Therefore, studying the evolution of bacterial operons provides valuable insight into understanding the biological roles of genes involved in environmental adaptation. However, no systematic approach has yet been devised to examine both the evolutionary relationships of operon encoded genes and the evolution of operon organization as a whole. To address this challenge, we present a novel method to study operon evolution by integrating phylogenetic tree based clustering and genomic-context networks. We apply this approach to perform the first systematic survey of all known synthase dependent bacterial biofilm machineries, demonstrating the generalizability of our approach for operons of diverse size, protein family composition, and species distribution. Our approach is able to identify distinct biofilm operon clades across phylogenetically diverse bacteria, resulting from gene rearrangement, duplication, loss, fusion, and horizontal gene transfer. We also elucidate different evolutionary trajectories of Gram-negative and Gram-positive biofilm production machineries, and in a companion paper (BIORXIV/2019/768473) present the experimental validation of a novel mode of biofilm production in a subset of Gram-positive bacteria.

## INTRODUCTION

The generation of novel genomes through next generation sequencing is creating a wealth of opportunities for understanding the evolution of biological systems. A key challenge is the development of robust and systematic approaches that allow genes not only to be classified into functional categories, but also infer evolutionary relationships. In bacterial genomes, functionally-related genes corresponding to metabolic pathways or protein complexes are often encoded by neighbouring co-transcribed loci, termed an operon. Computational prediction of operons based on the conservation of short inter-genetic distances has frequently been used to assign functions to uncharacterized genes from the known biological roles of their co-conserved neighbours(1–3). Analyzing patterns of sequence divergence within each gene can subsequently yield insights into species-specific functionalities. However, genes in an operon do not function isolation but tend to form parts of higher-order, biological modules (e.g. protein complexes or metabolic pathways). Consequently, analysing evolutionary events in an operonic context provides additional opportunities to better infer functional relationships. For example, while sequence divergence has the potential to impact the function of a single gene, evolutionary events that alter operon structure (e.g. rearrangements, duplications, gains and losses) have the potential to dramatically alter the overall function of the operon(4,5).

Due to the lack of a systematic framework, very few studies have attempted to examine the role of evolutionary events on operon structure(6,7). Phylogenetic-tree based classification of 197 ATP binding motif sequences associated with operon-encoded bacterial ATP-binding cassette (ABC) transporters was successful in resolving two evolutionarily distinct transporter clades associated with import and export functions(8). Gene duplications have been shown to play an important role in driving protein superfamily expansion among bacterial genomes and its frequency is significantly correlated with genome size(9). The study of co-localized “gene blocks” across bacteria has also shown that gene duplication, loss, and rearrangement play important roles in shaping the large-scale organization of bacterial genomes(7). Key to these analyses is the use of a rigorous and systematic approach for assigning genes into evolutionarily related ‘families’ that are likely to share similar functions. However, the inference of biological function based on sequence similarities of genes or proteins are often complicated by functional divergence arising through recent gene duplication events. A variety of metrics have been employed for determining the relatedness of genes and their protein products from which groups (i.e. clustering) can be defined. These metrics include: evolutionary distances derived through the construction of phylogenetic trees(10–13); global protein sequence similarities(14–16); and shared sequence features such as conserved amino acids at specific sites or shared amino acid subsequences(17,18). The aim of these approaches is to automatically resolve large protein families comprising potentially thousands of genes into a smaller number of clusters defining evolutionarily related subfamilies with similar biological roles. Here we build on these methods and present a novel framework for the systematic classification and analysis of genes in the context of operons. Focusing on synthase-dependent exopolysaccharide (EPS) biosynthetic machineries, we use our framework to explore how gene divergence in combination with duplication, loss, and rearrangement events have shaped the evolution of EPS operons, and may have influenced the biofilm producing capabilities of evolutionarily diverse bacteria.

EPS are an important constituent of bacterial biofilms that not only ensure survival in response to limited nutrient availability, but are also involved in antibiotic tolerance, immune evasion and serve as virulence factors in many clinically relevant pathogens (19–21). Distinct mechanisms have been identified in the production of bacterial EPS, including the well-characterized Wzx/Wzy and ABC transporter-dependent pathways involved in capsular polysaccharide and lipopolysaccharide secretion in Gram-negatives (22), and the more recently identified synthase-dependent systems(23).

Typically, synthase-dependent EPS systems are organized as discrete operons comprised of genes encoding: 1) an inner membrane associated polysaccharide synthase and copolymerase subunit; 2) a regulatory domain responsible for binding the intracellular signaling molecule cyclic-3’,5’-dimeric guanosine monophosphate (c-di-GMP); 3) periplasmic polysaccharide modification enzymes; and 4) a periplasmic tetratricopeptide repeat (TPR) domain coupled with an outer membrane pore(23). This operonic organization allows bacteria to acquire complete EPS functionality through discrete lateral gene transfer events and may act as a key driver in niche adaptation(24). To date five synthase-dependent EPS have been identified: cellulose, acetylated-cellulose, poly-β-1,6-*N*-acetyl-D-glucosamine (PNAG), alginate and the Pel polysaccharide. While much interest has focused on the molecular basis of biofilm formation, much less is known about how these systems have propagated across bacterial taxa. Further, it is not known how EPS operons evolve to help bacteria adapt to diverse environments and, from a human health perspective, establish infection and cause disease. While a previous survey of cellulose EPS machineries has been reported(25), a comprehensive systematic analysis of all EPS machineries is lacking.

In this study, we describe a phylogenetic tree-based clustering method for defining protein sequence subfamilies and apply it to study the evolutionary relationships of operons. This approach was employed for the systematic classification of EPS operons predicted from a survey of over a thousand bacterial genomes. Applying a graphical visualization approach, we demonstrate that phylogenetic clustering enables the resolution of discrete EPS operon clades, enabling the identification of distinct operon organizations across diverse bacterial phyla which have been shaped by locus duplications, losses, and rearrangement events. For example, we demonstrate the biological implications of operon evolution that has been shaped by horizontal gene transfer (HGT) and subsequent divergence, for two cellulose operon clades among Proteobacteria which correspond to the production of cellulose polymers with different structural organizations. Although Pel production was initially identified and characterized in *Pseudomonas aeruginosa*(26), our approach also identified a number of *pel*-like operons in some *Bacillus* spp. and other Gram-positives. A subset of these systems appear to be regulated by the intracellular signalling molecule c-di-GMP. In our companion paper (BIORXIV/2019/768473) we experimentally validate these finding and demonstrate the production of Pel by the Gram-positive *Bacillus cereus* ATCC 10987 that is regulated by c-di-GMP.

## RESULTS

### A Systematic Survey of Bacterial EPS Operons Reveals EPS Systems Across Bacteria of Diverse Lifestyles and Environmental Niches

To examine the phylogenetic distribution of EPS systems, we first performed a systematic survey of all five previously characterized synthase-dependent EPS systems (cellulose, acetylated-cellulose, PNAG, Pel, and alginate) (**Supplemental Tables 1 & 2**) through an iterative hidden Markov-model (HMM) - based search strategy and subsequent genomic-proximity based reconstruction of 1861 complete reference and representative bacterial genomes (downloaded April 20, 2015 - see **Methods**). We identified 407 cellulose, 321 PNAG, 146 Pel, 64 alginate, and 4 acetylated-cellulose EPS “operons” defined as comprising at least: 1) a polysaccharide synthase subunit; and 2) one additional locus involved in EPS modification or transport as defined previously(19) (**Supplemental Table 3**). These could be allocated to 367, 288, 140, 60 and 4 different bacterial species, respectively (**Figure 1**). Each type of system was largely associated with proteobacteria, with cellulose, PNAG and Pel additionally featuring operons from Bacilli and Clostridia, which to our knowledge have not been previously reported. *pel* operons exhibited the greatest diversity of bacterial families (shannon index of bacterial families – 2.74) with representation from Thermotogales, Actinobacteridae and Rubrobacteridae, among others. PNAG was significantly enriched in pathogen genomes (161/289 - 56%; T-test p-value <0.005). Conversely, Pel (84/140 - 60%; T-test p-value < 0.001), alginate (39/60 - 65%; T-test p-value < 0.001) and cellulose (187/367 - 51%; T-test p-value < 0.001) were significantly enriched in non-pathogen genomes (**Figure 1 and Supplemental Figure 1**). Interestingly, both cellulose and PNAG operons were significantly associated with genomes with host-associated lifestyles (T-test p-values < 0.001). While most genomes contain only a single synthase-dependent EPS system, we observed many instances of co-occurrence, with cellulose and PNAG systems being the most common combination (83 genomes), followed by alginate and *pel* (20 genomes). Notably, all species possessing three systems were *Pseudomonas* spp., e.g.: *Pseudomonas protegens* strains Pf-5 and CHA0 (alginate, *pel* and PNAG); P*seudomonas fluorescens* SBW25 and *Pseudomonas* sp. TKP (acetylated-cellulose, alginate, and PNAG).

**Figure 1.**
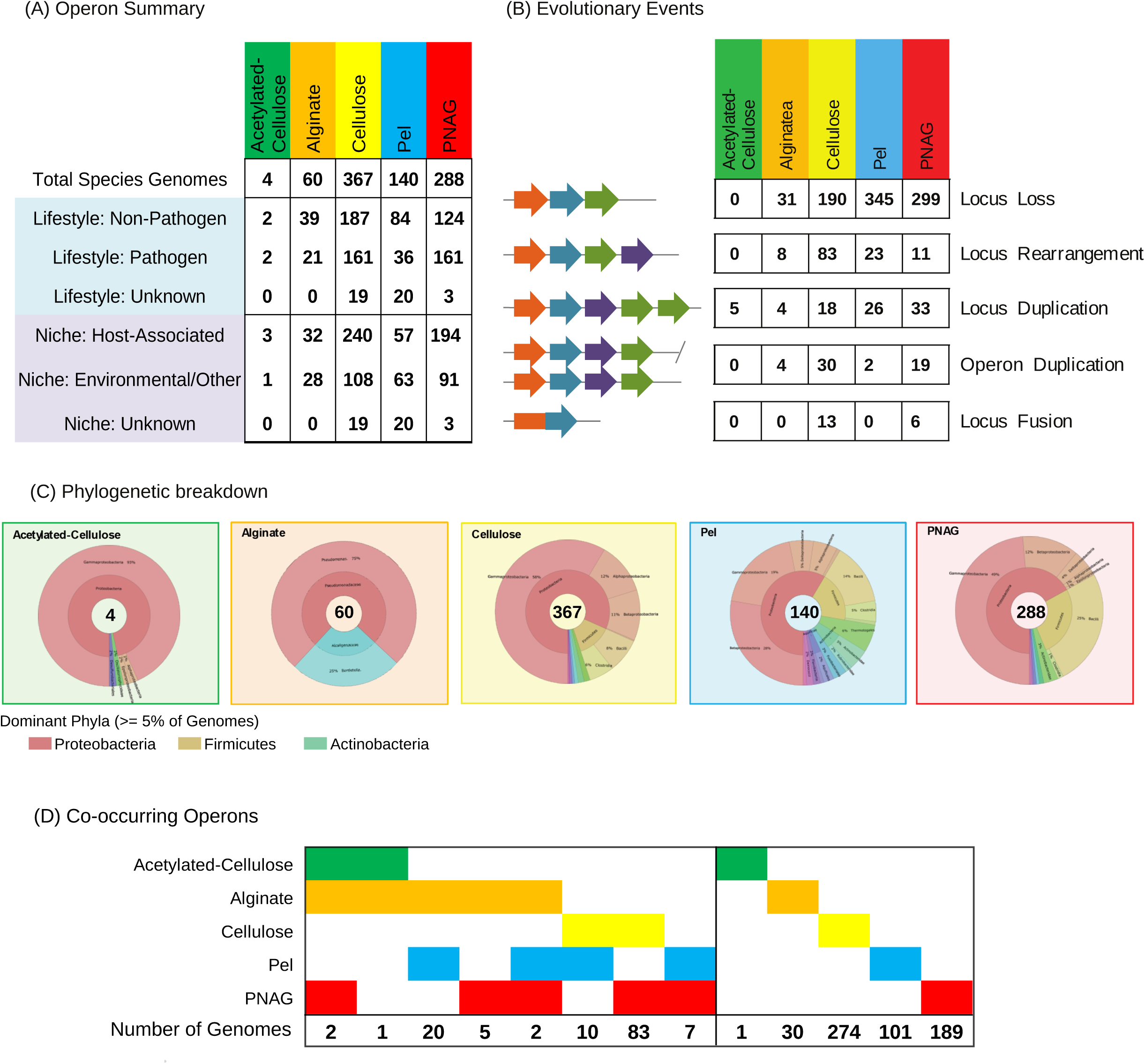
Summary of Predicted Bacterial EPS Operons. (A) Number of predicted EPS operons are summarized by bacterial lifestyle (pathogen, non-pathogen, unknown) and corresponding niche (host-associated, environmental/other, unknown). (B) Evolutionary events associated with EPS operons: Locus loss (core EPS operon loci not detected by HMM searches); Locus rearrangement (EPS operons featuring locus orderings that differ from the canonical operon for that type – **Supplemental Table 1**); Locus duplication (defined by two loci possessing a significant match to the same EPS HMM within the same operon); Operon duplication, defined as a genome encoding two copies of the same type of EPS system, separated by greater than 10 Kbp; Locus fusion, loci possessing significant matches to multiple EPS HMMs. (C) Phylogenetic distribution of EPS operons visualized by Krona(88). (D) Patterns of EPS operon co-occurrences indicating the frequency of specific operon combinations within a single genome.

### Evolution of EPS Operons is Driven by Gene Duplication, Loss and Rearrangements

The processes underlying EPS operon evolution across diverse bacterial phyla is poorly understood.. We examined how operon organization is influenced by the following evolutionary events that are likely to affect EPS production capabilities among bacteria: 1) single locus or whole operon duplications, corresponding to dosage effects altering the level of EPS modification or export; 2) locus losses, that may indicate a reduction or loss in EPS production or modification, or may suggest supplementation of the lost function with a novel gene; 3) operon rearrangements which may affect the regulation of EPS production through the order of expression of individual EPS system components; and, 4) gene-fusions, resulting in enhanced co-expression of interacting subunits.

For each set of predicted EPS operons, the resulting number of operon evolutionary events were assessed relative to the locus composition and ordering of reference Gram-negative experimentally characterized operons defined from previously published studies(19,28–32). With the exception of acetylated-cellulose, locus losses were found to be the most frequent event (∼46% of predicted operons lacked one or more reference loci), and occurred with the greatest frequency for Pel which exhibited an average loss of 2.6 loci lost per operon (**Supplemental Table 4**). Among all EPS systems the majority of locus losses were associated with the outer-membrane pore encoding loci (441 / 993 - 44% of all locus loss events identified) among Gram-positive species (**Supplemental Table 4**), consistent with the lack of an outer-membrane bilayer in Gram-positive membrane architectures. Operon rearrangements were the next most frequent evolutionary events (∼ 39%), largely associated with cellulose operons(25) (327 / 407 – 80%). Focusing on duplication events, within-operon loci duplications tended to be more common than whole-operon duplications (2 or more core EPS loci identified >= 1 Kbp apart), with the exception of cellulose operons (29 whole operon duplications v 24 loci duplications). All duplicated operons were found to be separated by at least 10 kbp, suggesting they may have been acquired through HGT rather than tandem duplication of a pre-existing operon(32,33).

### Systematic Phylogenetic Distance-Based Clustering of EPS Operon Loci and Genomic-Proximity Networks Identifies Evolutionarily Distinct Operon Clades

To better understand how these evolutionary events may have altered operon function, we next devised an agnostic, systematic classification strategy to cluster each family of EPS operon loci on the basis of phylogenetic distance (**Figure 2A; see Methods**). In brief, for each EPS operon locus, multiple sequence alignments were generated and used to construct phylogenetic trees. From these trees, we defined sets of clusters through an iterative scan of the tree structure that captures an increasing sequence distance between family members, starting at the leaves and ending at the root. During this scan, sequences are grouped into a cluster if they share a common node (i.e. are within a specified evolutionary distance). To define the optimal set of clusters for each locus, we then applied three cluster quality scoring schemes (Q1, Q2 and Q3) based on the following metrics: proportion of sequences clustered (to maximize the number of sequences clustered); average silhouette score (to minimize the occurrence of clusters containing highly divergent sequences); and the Dunn index (to maximize the separation of closely related sequences from divergent sequences). For each scoring scheme, we defined the optimal pattern of clustering based on the evolutionary distance (expected number of substitutions per site) derived from a maximum-likelihood constructed phylogenetic tree (**see Methods for more details)** that maximizes the quality score. Comparisons across scoring schemes (see below) for cellulose operon loci identified Q2 as providing the most informative sets of clusters. Applying this scoring scheme to all EPS loci revealed the average number of sequence clusters generated correlated with the total number of operons predicted for each type of EPS system (**Figure 2B**), which further corresponded to the underlying differences in species distributions of EPS systems (**Figure 1C**). For example, the cellulose system was predicted to have the largest average number of sequence clusters overall (30 clusters) and also had the greatest species diversity (shannon index 2.16 – **Supplemental Figure 2**) compared to all other systems. Furthermore, for each EPS system the variability of the number of sequence clusters predicted per locus (**Figure 2C**) suggests differing degrees of locus evolution that are likely to be the result of different structural and functional constraints. For example, a higher degree of conservation would be expected for glycosyl transferase (GT) subunits to maintain efficient co-ordination between polymerization and inner-membrane transport of EPS, while increased variability of periplasmic modification enzymes suggests that only a subset of highly conserved motifs are required to carry out polysaccharide modification reactions.

**Figure 2.**
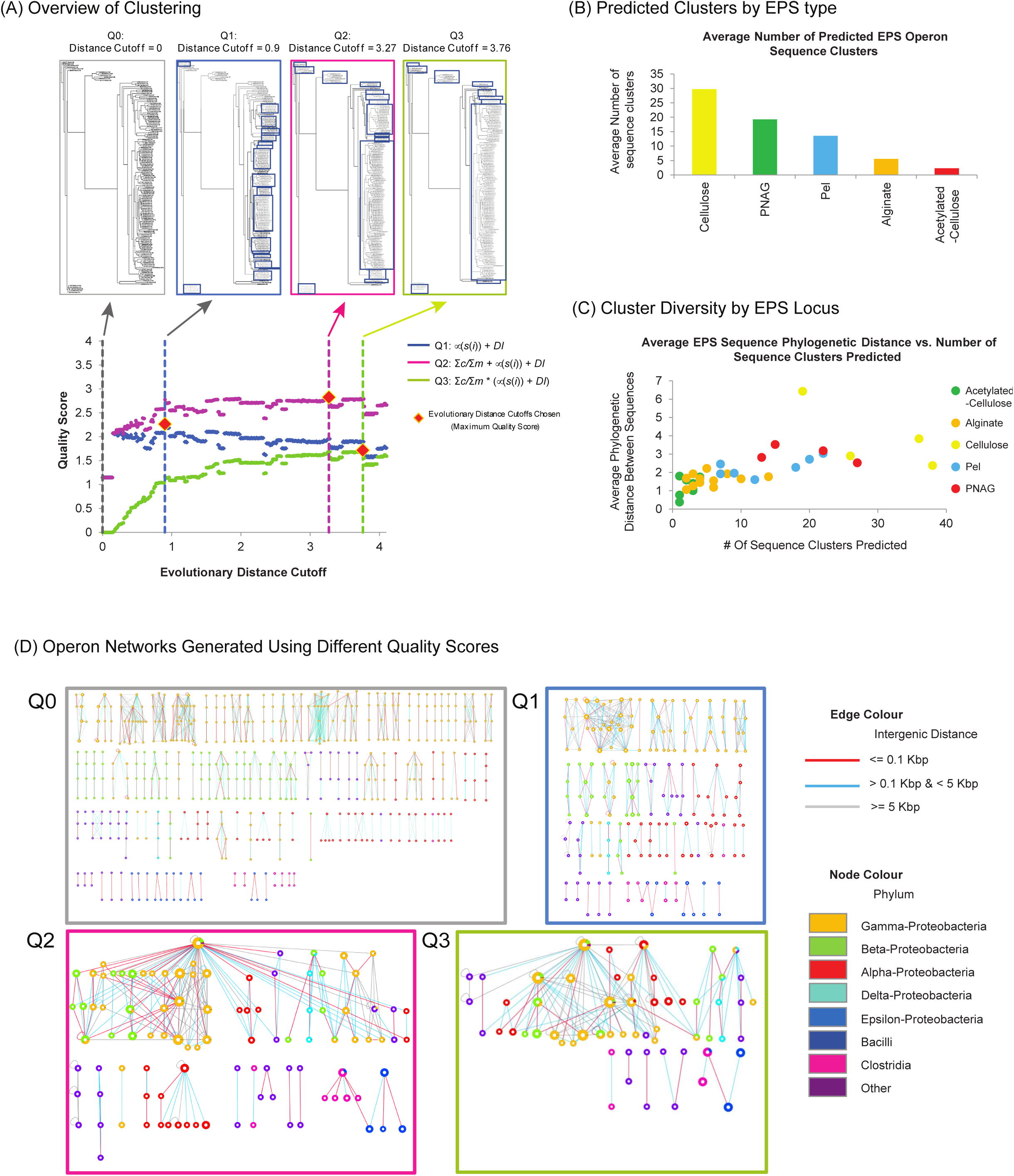
Clustering of EPS Loci. (A) Schematic illustrating the process of scanning through a phylogentic tree and identifying sets of clusters associated at different evolutionary distance cutoffs. Here evolutionary distance is defined as the number of expected amino-acid substitutions normalized over the multiple sequence alignment length. To identify optimal patterns of clusters, we examined three scoring schemes (Q1, Q2 and Q3). Q1 is defined as the sum of the average silhouette score for all clusters: μ(s(*i*)), and the Dunn index (DI). Q2 is defined as the sum of the proportion of sequences identified in clusters (Σ*c*/Σ*m*), μ(s(*i*)) and *DI*. Q3 is defined as the product of Σ*c*/Σ*m* and the sum of μ(s(*i*)) and DI. For the family of genes related to the *bcsA* locus, each scoring scheme identifies a different optimal evolutionary distance cutoff resulting in defining different sets of clusters. (B) Graph illustrating the average number of sequence clusters predicted (sum of # of clusters over all loci / total number of EPS loci) for each type of EPS operon. (C) Graph illustrating the average evolutionary distance of EPS loci cluster members with other members of the same cluster. (D) Cellulose operon networks generated using the different types of scoring scheme cutoffs used in (A). For each network, nodes indicate clusters of sequences representing individual cellulose loci, edges indicate genome proximity between the two linked loci. Nodes are organized into sets of four, ordered from top to bottom as *bcsA, bcsB, bcsZ* and *bcsC*. Node size indicates the number of family members associated with that locus cluster. Node colour indicates phylogenetic representation of cluster members. Edge colour indicates genomic proximity of phylogenetic clusters. At higher evolutionary distances (as defined by Q2 and Q3), networks yield more informative patterns of evolutionary relationships as illustrated by larger clusters of loci featuring larger number of interconnections.

To compare patterns of clusters identified by each scoring scheme, we applied the three scoring schemes to each set of genomically-neighbouring protein sequences assigned by HMM searches to a given cellulose locus (*bcsA, bcsB, bcsZ*, and *bcsC*). From the resulting clusters we generated operon genomic-proximity networks (**Figure 2D**). These networks provide a visual display of the conservation of individual loci, together with their respective genomic proximity to yield patterns of sequence divergence associated with the emergence of distinct forms of operon organization. In the absence of any clustering (Q0), the network trivially resolves into individual operons featuring up to four loci. Applying the Q1 scoring scheme to each locus, the network reveals a variable number of clusters across operon loci, with each cluster generally comprising sequences belonging to the same bacterial genus. Application of the Q2 scoring scheme results in the generation of clusters of increased size, encompassing species featuring distinct operon organizations and compositions. For example, two distinct lineages of alpha-proteobacterial cellulose operons can be easily distinguished, one of which is more closely related in sequence and composition to gamma-proteobacterial operons, and a second which lacks two loci and appears evolutionarily divergent from gamma-proteobacterial operons(25). However, these distinctions were more difficult to resolve using the Q3 scoring scheme due to clustering of highly divergent sequences. Given the trade-off between clustering highly divergent sequences (Q3) with the depiction of individual operons (Q1), we applied the Q2 scoring scheme to generate clusters for all EPS loci (**Supplemental Table 5**).

Using this locus-specific phylogenetic clustering approach, we were able to devise a classification scheme to define EPS locus clades based on the average evolutionary distance of a group of clustered locus sequences to a reference operon sequence (**Supplemental Table 3**). For example, the cellulose polysaccharide synthase locus, *bcsA*, from *Escherichia coli* is assigned to clade 1, while divergent alpha-proteobacterial species including *Rhodobacter sphaeroides* are assigned to clade 2. We further resolved operons into distinct groups based on the genomic co-occurrence patterns of locus clades; e.g. for the cellulose operon (*bcsABZC*) we identify clade combinations of 1:1:1:1, 1:2:2:2 and 1:3:5:3, which correspond to operons identified in *Escherichia* spp. and other closely related enterobacteria, *Klebsiella* spp., and *Burkholderia* spp., respectively.

### Phylogenetic Clustering and Genomic Proximity Networks Reveal Evolutionary Events Driving EPS Operon Divergence

Focusing on cellulose EPS operons, relative to BcsA, the polysaccharide synthase subunit, the three other subunits (BcsB, BcsZ and BcsC) display greater sequence diversity as indicated by a larger number of sequence clusters (**Figure 3**). Detailed structure-function studies of the BcsA-BcsB inner-membrane cellulose synthase complex, outlined below, illustrate how these findings are consistent with their known functional roles. Further inspection of the cellulose operon network identifies a number of sub-networks comprised of taxon-specific loci clusters associated with distinct patterns of operon organization as illustrated through the following examples: 1) a subnetwork comprised of loci from several beta-proteobacteria, represented here by *Burkholderia cenocepacia* and *Pandoraea* promenusa (**Figure 3(i)**), which feature a rearrangement of the *bcsA* locus and novel locus gains (also supported by inspection of corresponding Genbank genomic annotations) as indicated by a genomic distance of > 0.1 Kbp between *bcsA* and the neighbouring locus *bcsC*; 2) a subnetwork composed of loci from several species of the alpha-proteobacterial *Zymomonas*, feature rearrangement of *bcsZ* and/or the loss of *bcsB* or *bcsZ*; further inspection reveal such losses to be due to gene fusion events (**Figure 3(ii)**); 3) a subnetwork composed of loci from a separate group of alpha-proteobacteria, reveals a diverse set of *bcsB* loci that additionally lack the *bcsC* outer membrane pore (**Figure 3(iii)**); and 4) a subnetwork of loci from a group of gamma-proteobacteria reveal instances of HGT and divergence (**Figure 4A**). In this latter example, our network identifies two distinct clades of operons, sharing a common group of *bcsA* loci, but featuring two evolutionarily divergent sets of *bcsB, bcsZ* and *bcsC* loci which co-occur in several genomes separated by inter-genic distances greater than 10kBp. Detailed investigation of the operonic arrangements of species possessing single copies of either of these clades of operons, reveal two distinct loci organizations: the first representing the canonical cellulose locus order (clade A1), *bcsABZC*, found among *Escherichia coli* and *Salmonella enterica* strains; the second represents a non-canonical locus ordering (clade B1), in which the periplasmic glycoside hydrolase, BcsZ, has undergone a rearrangement, *bcsABCZ*, and is found among *Dickeya, Erwinia* and *Pantoea* spp. (**Figure 4B**). Of note, we found that several species (e.g. *Enterobacter* and *Klebsiella* spp.) possess both operon clades, which have previously been inferred as originating by HGT(19) and is further supported by our phylogenetic clustering assignments (**Figure 4C**). Furthermore, we identified two additional divergent BcsB sequences associated with a novel organization of operon clade B1 and include several loci with other roles in cellulose production (designated operon clade B2; **Figure 4D**). The divergence of BcsB sequences associated with clade B2 were also found to distinguish bacterial genomes possessing multiple cellulose operons of distinct evolutionary lineages: *Proteus mirabilis* (2 cellulose operons: Clades A1 and B2) and *Enterobacter* spp. (3 cellulose operons: Clades A1, B1 and B3) (Figure 4E). Additional sequence database searches revealed that the non-core loci associated with operon clades B2 and B3 share functionally homologous loci to the cellulose accessory protein D (AxcesD), which has been characterized as increasing the efficiency of cellulose production in the *Acetobacter xylinus* cellulose synthase complex(34); GalU an uridine triphosphate (UTP)-glucose-1-phosphate uridylyltransferase involved in cellulose precursor biosynthesis; and an additional uncharacterized locus predicted to possess both PAS_9 and GGDEF signalling domains, indicating the potential adaptation in *Proteus* and *Enterobacter* spp. to produce varied forms of cellulose upon different environmental stimuli(35).

**Figure 3.**
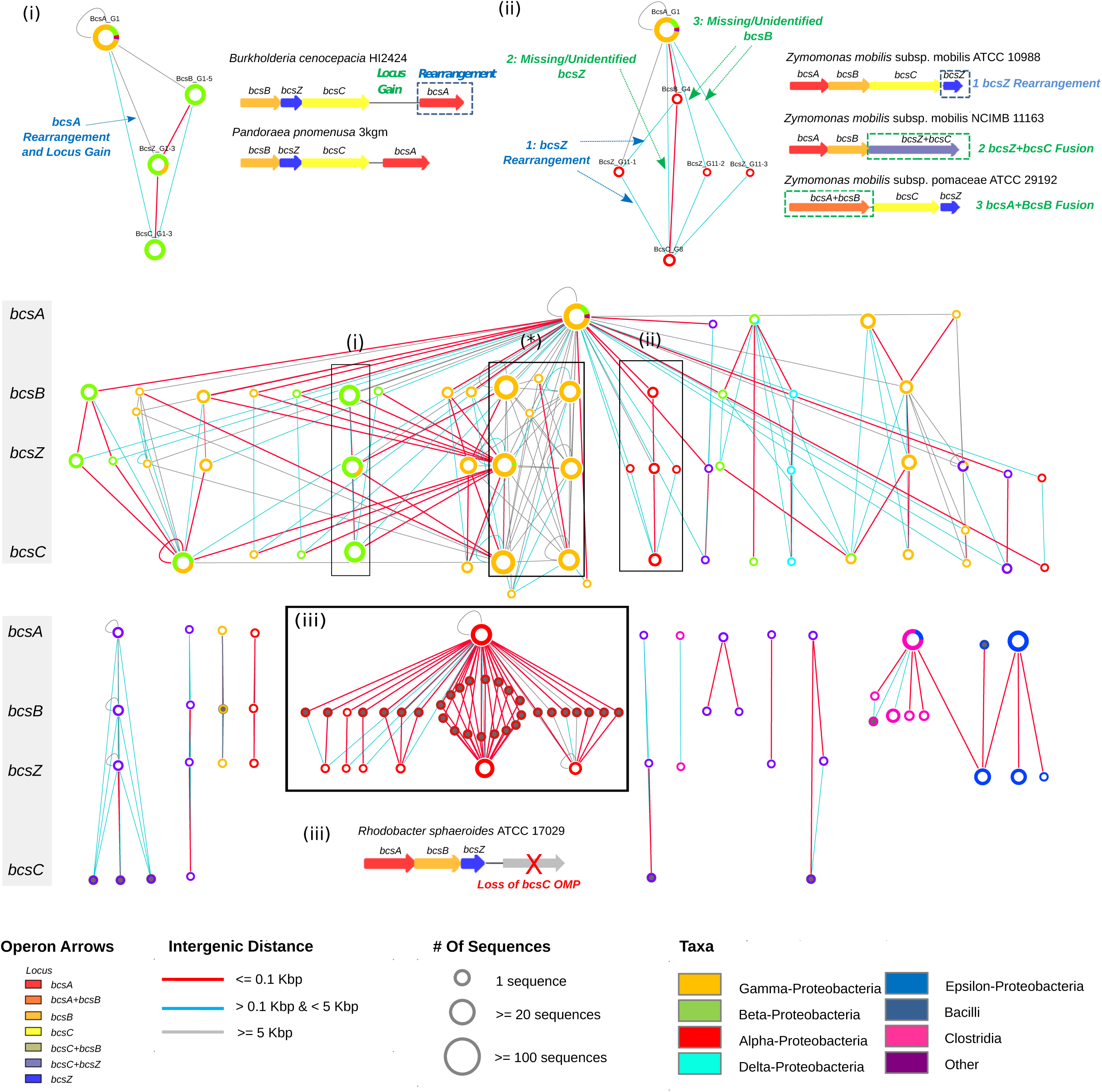
Genomic-Proximity Network of Phylogenetically Clustered Cellulose Operons. Phylogenetically clustered operon loci are arranged vertically with respect to the canonical ordering of the cellulose operon (indicated by grey side bar). Inset boxes depict selected examples of cellulose operon clades, illustrating how the network can inform on evolutionary events: (i) Rearrangement of *bcsA* among betaproteobacteria – Here, *bcsA* appears in closer proximity to *bcsC* than to *bcsB* or *bcsZ* (as indicated by a cyan coloured edge for the former and a grey coloured edge for the latter). Further the cyan edge indicates a relatively large intergenic distance, suggesting a locus gain between *bcsA* and *bcsC*, confirmed upon inspection of the genome of *Burkholderia cenocepacia*; (ii) Rearrangement and gene fusions in alphaproteobacterial – in examples 1 and 2, the red edge indicates operons in which *bcsB* is closer to *bcsC* than *bcsZ*, the cyan edges suggest that *bcsZ* is present, but appears after *bcsC* (example 1), while in other operons, *bcsZ* appears missing (example 2). Detailed inspection of example operons (e.g. *Zymomonas* spp.) reveals the fusion of the periplasmic hydrolase and outer membrane pore (BcsZC), in example 3, the apparent loss of *bcsB* in another *Zymomonas* spp. is explained by a fusion between the inner membrane cellulose synthase complex subunits (BcsAB); (iii) Loss of outer membrane pore, BcsC, and divergence of the inner membrane cellulose co-polymerase, BcsB, in alphaproteobacteria – in these taxa, BcsB appears highly divergent (as indicated by their identification through more sensitive HMM searches – grey nodes) and no BcsC was identified (confirmed through inspection of representative operons). Further interpretation of the operons identified in the box denoted with a ‘*’, which represent HGT events, are illustrated in Figure 4. Node size indicates the relative number of sequences per phylogenetic cluster; node colouring represents the taxonomic distribution of loci for a given cluster; edges connect clusters which co-occur in the same genome(s); edge colour indicates the genomic-proximity of loci clusters. The network was visualized using Cytoscape 3.5.1(87).

**Figure 4.**
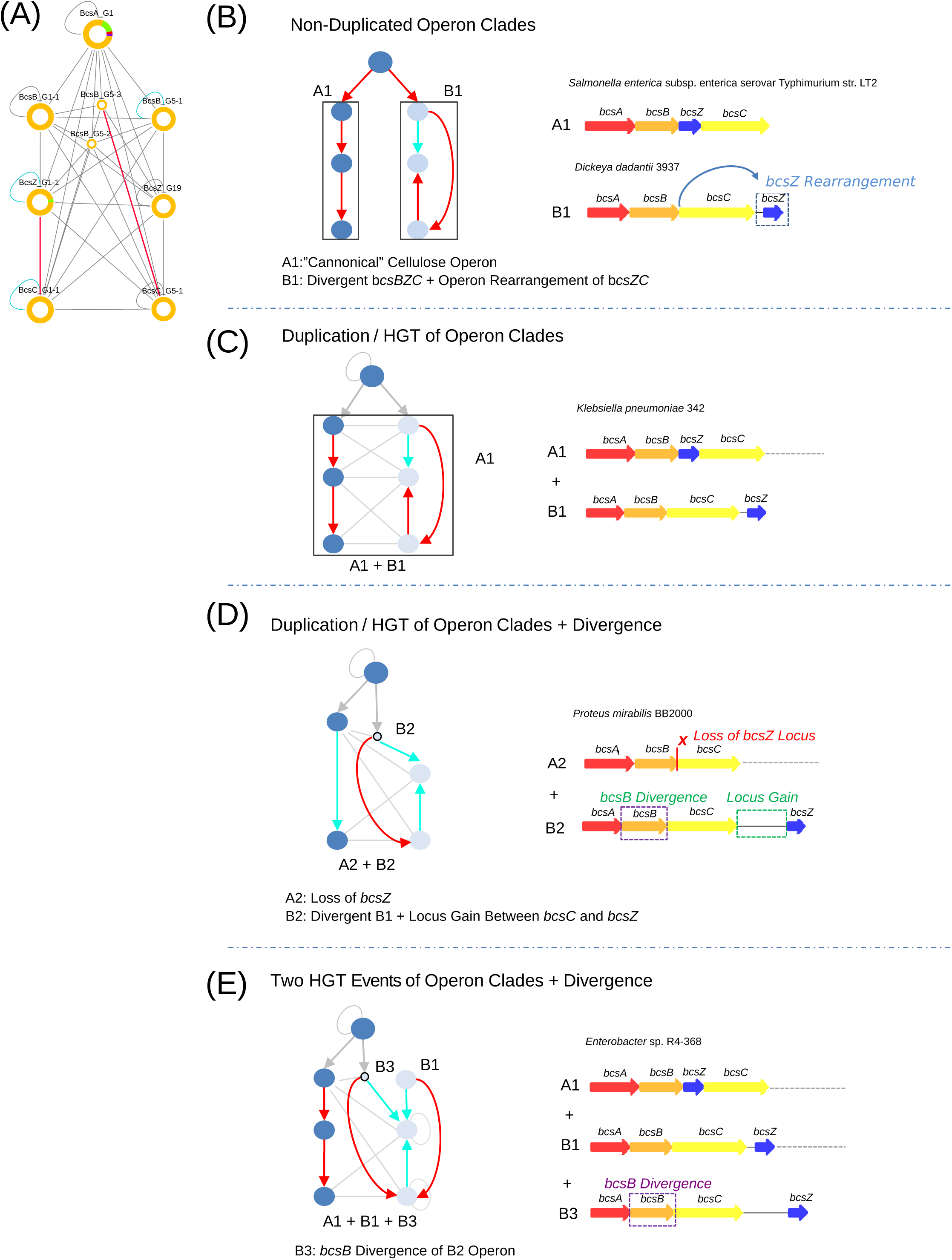
Horizontal Gene Transfer of Cellulose Operons Identified From Analysis of the Genomic-Proximity Network. Here we show how a subgraph (A) from the global cellulose EPS operon genomic-proximity network (**Figure 3(*)**), may be interpreted to reveal HGT events involving two distinct gamma proteobacterial operon clades, A (canonical *bcsABZC*) and B (*bcsABC-Z*). (B) Examples of operons in two species which possess either a single A1 (“canonical”) or B1 (rearrangement of *bcsZC*) operon clade. (C) Example from *Klebsiella pneumoniae* in which a single genome contains both A1 and B1 operons, indicating a HGT event. (D) Example from *Proteus mirabilis* featuring two copies (designated A2 and B2 respectively) of the cellulose EPS operon, which appear to be divergent forms of A1 and B1: A2 features an apparent loss of the *bcsZ* locus from A1; B2 features a locus gain between *bcsC* and *bcsZ* from B1. Example from *Enterobacter* spp. in which the genome carries three copies of the cellulose EPS operon. In addition to clade A1 and B1 operon arrangements, a further operon (designated B3) appears in which *bcsB* has diverged from a B2 clade operon. Arrows within the network schematics depict the order of loci within the operon and are coloured according to intergenic distance: red < 100bp; cyan >100bp & <5 Kbp; grey >5Kbp.

### Genomic-Proximity Networks of *pel* Operons Reveal a Novel *pel* Locus in the Gram Positive Bacterium, *Bacillus cereus* that is Regulated by c-di-GMP

Examination of the genomic-proximity networks of *pel* loci also reveal novel operon organizations across phylogenetically divergent bacteria (**Figure 5**). As with cellulose loci *bcsA* and *bcsZ*, we identify examples of operon rearrangements involving *pelB* (outer membrane transport pore + TPR domain) loci and *pelA* (periplasmic modification hydrolase) (**Figure 5(ii), (iiib), (iv)**), across several species associated with diverse environments. Again consistent with our findings for cellulose, we noted loci losses and acquisitions. Although it has not been demonstrated that the pel operon forms a trans-envelope biosynthetic complex, the ordering of operon loci has been shown to play an important role in the assembly of macromolecular complexes(36) and optimizing biosynthetic pathways(37), suggesting that there exists a functional coupling between pel outer-membrane transport and periplasmic modification(38). However, the effects of these rearrangement events on Pel production still remain to be experimentally investigated.

**Figure 5.**
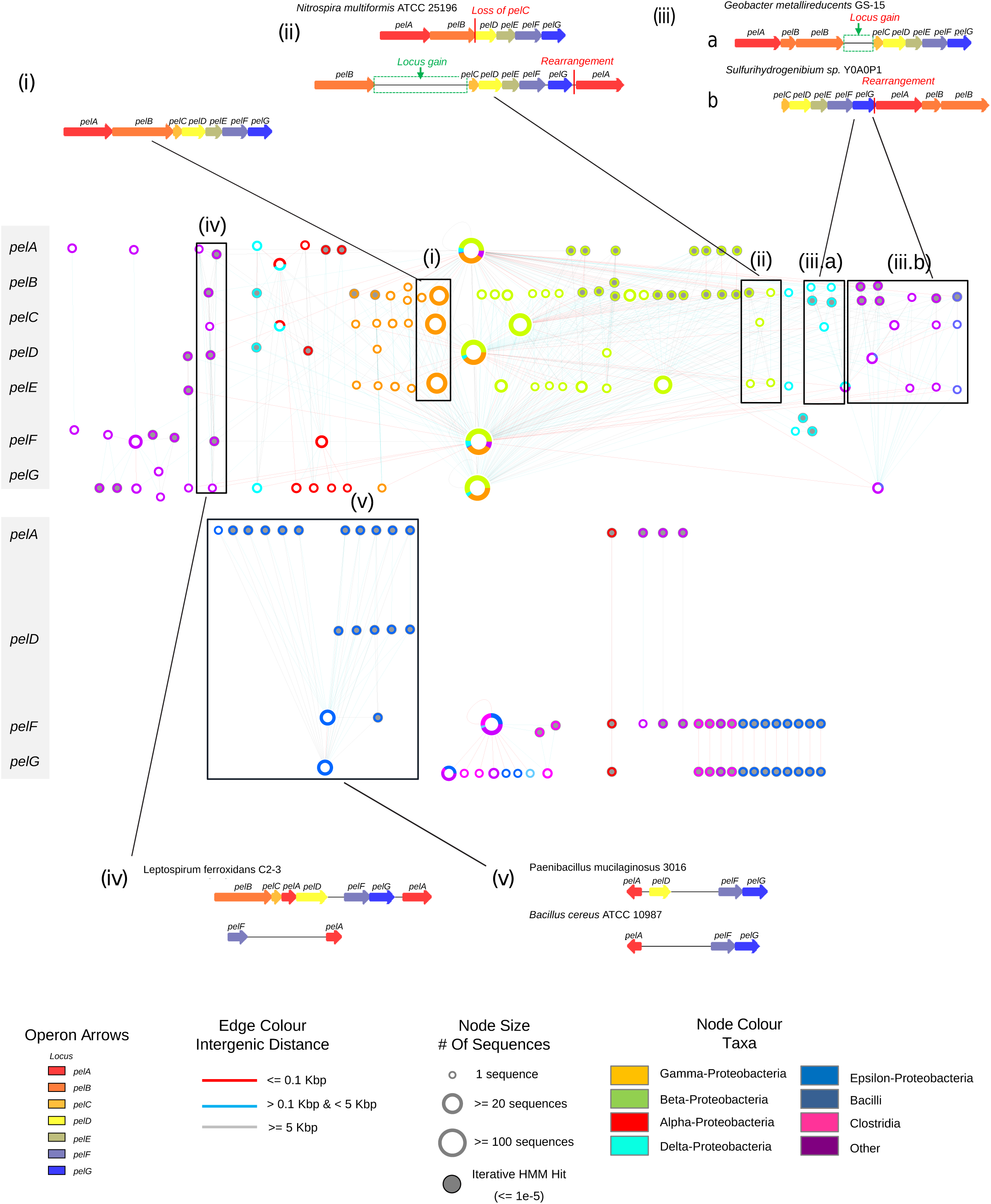
Genomic-Proximity Network of Phylogenetically Clustered *pel* Operons. Phylogenetically clustered operon loci are arranged vertically with respect to the canonical ordering of the *pel* operon (indicated by grey side bar). As for Figure 4, inset boxes depict selected examples of *pel* operon clades, illustrating how the network can inform on evolutionary events: (i) Canonical organization of the *pel* operon, as defined in the *Pseudomonas aeruginosa* genome.; (ii) Duplication of the pel operon in *Nitrosospira multiformis* with subsequent evolution through locus gain and loss, as well as rearrangement of *pelA*; (iii) *pelB* fission, locus gain and rearrangement in aquatic thermophilic species; (iv) A potentially novel duplicated pel operon identified in *Leptospirillum ferrooxidans* comprised of divergent *pelA* and *pelF* loci; (v) *pel* operons identified in Gram-positive species including divergent *pelD* loci involved in regulation through cyclic di-GMP. Node size indicates the relative number of sequences per phylogenetic cluster; node border colouring represents the taxonomic distribution of loci for a given cluster; grey filled nodes indicate loci predicted by iterative HMM searches; edges connect clusters which co-occur in the same genome(s); edge colour indicates the genomic-proximity of loci clusters. The network was visualized using Cytoscape 3.5.1(87).

We also observed a high degree of overall conservation among components which are known to play key roles in Pel biogenesis, such as the putative polysaccharide synthase (PelF), putative inner-membrane protein (PelG), hydrolase/deacetylase (PelA) and cyclic-di-GMP receptor (PelD)(21). In contrast, a greater degree of divergence can be seen among inner (PelE) and outer-membrane (PelB, PelC) transport associated loci, which appear to follow a consistent pattern of clustering across bacterial phyla suggesting co-evolution of potentially physically interacting components, however no evidence of interaction has been shown to date.

Our genomic proximity network revealed two distinct clades comprising several Gram-positive species (**Figure 5(v)**). Of the synthase dependent EPS operons known to date, only PNAG production has been genetically and structurally characterized in Gram-positive Staphylococci(39). Operons reconstructed from initial HMM searches identified putative *pel* operons in several Gram-positive bacteria, comprised of the GT encoding PelF and the PelG putative transport protein (**Figure 5**). To determine whether these were bona-fide *pel* operons with additional loci, iterative HMM searches were performed including additional protein sequences from predicted pel operons, revealing additional loci including a homolog of PelD (**Supplemental Figure 3**). C-di-GMP signaling in Gram-positive bacteria is less well characterized(40) and this finding provides evidence for its role in regulating biofilm formation in these species. In our companion paper, we have experimentally validated our predictions by showing that single deletion knockouts of the predicted *B. cereus* operon loci result in a loss of EPS production, and that PelD regulates EPS production through binding of c-di-GMP (BIORXIV/2019/768473).

### Genomic-Proximity Networks of PNAG Uncover Locus Loss and Duplication Events in Pathogenic and Environmental Bacteria

To examine how locus duplication, loss, and rearrangement events have contributed to the evolution of PNAG operons across bacterial phyla, selected examples of *pga* operon clusters were identified and compared (**Supplemental Figure 4**). For example, within a group of enterobacteria possessing related *pgaD* loci, there exist a number of closely related pathogen enterobacteria that have lost *pgaA* (*E. coli* ETEC H10407), as well as *pgaB* (*Shigella flexneri* 5 str. 8401), suggesting the recent loss of the ability to produce PNAG (**Supplemental Figure 4(i**.**a)**); in the case of *S. flexneri* this loss may be due to adaptation to an intracellular mode of infection(41). Similarly, no *pga* operons were detected among *Salmonella* spp. genomes surveyed in this study, consistent with the loss of PNAG production previously associated with an intracellular pathogenic lifestyle(42).

Based on the divergence of *pgaB* loci, we also identified *pga* operon clades corresponding to partial and whole operon duplications in aquatic bacteria, including a partial duplication of the *pga* operon specific to *Acinetobacter baumannii* spp. and *Methylovora versatilis* 301, respectively (**Supplemental Figure 4(ii)**). Also, in environmental bacteria we discovered a novel *pga* organization resulting from rearrangement of *pgaC*, and a lack of *pgaB* and *pgaD* loci, which may have been too divergent to detect from inital HMM searches (**Supplemental Figure 4(iii)**). Although our HMM models were based on solely Gram-negative *pga* operon protein sequences, we also identified a number of Gram-positive *pga* operons consisting of *pgaB* and *pgaC* (**Supplemental Figure 4(i**.**b, i**.**c)**). Upon closer inspection these loci were found to correspond to *Staphylococcus* intercellular adhesion (ICA) loci *icaB* and *icaA*, respectively, suggesting a potential common evolutionary origin of synthase-dependent PNAG production between Gram-positive and -negative organisms.

A clade of *pga* operons were also identified possessing varying numbers of divergent *pgaC* loci resulting from repeated tandem duplication events (**Supplemental Figure 4(v)**). Despite lacking a detectable *pgaA* locus, a possible role of these gene clusters in EPS production was investigated. One member of this operon clade, *Thauera* sp. MZ1T, inhabits a wide range of environments, and is an abundant producer of EPS responsible for viscous bulking in activated sludge wastewater treatment processes(43). Furthermore, a recent mutagenesis study(44) demonstrated that biofilm-formation defective *Thaurea* mutants could be rescued by the complementation of the predicted *pgaB* deacetylase locus identified in the present study. Combined with our evolutionary clustering results, these findings suggest that the divergence of deacetylase and duplication of GT related-loci in PNAG biosynthesis have resulted in the emergence of a distinct operon lineage.

### Genomic Proximity Networks of Alginate Uncover Distinct Operon Clades in *Pseudomonas* spp. and Atypical Operon Architectures in Environmental Bacteria

Although the majority of alginate operons were predicted largely among *Pseudomonas* spp. genomes (**Supplemental Figure 5**), phylogenetic clustering and genomic-proximity network reconstruction revealed an array of events influencing alginate operon evolution. For example, two distinct alginate operon clades were identified among *Pseudomonas* spp., defined by whole operon duplication and rearrangement of alginate polysaccharide modification loci (**Supplemental Figure 5(i) and (ii)**). Also identified were divergent, “atypical”, alginate operons (**Supplemental Figure 5(iii)**) comprising extensive rearrangements and also losses of functionally related subsets of alginate loci, e.g. outer-membrane transport loci (*algKE*), and polysaccharide modification machinery (*algGXLIJF*). Closer examination of the alginate genomic-proximity network also indicated a greater number of clusters for *alg44* and a*lgX* loci, which were reflective of increased divergence among distinct alginate operon clades. Given that both loci play related roles in the regulation, polymer-modification, and assembly of the alginate EPS secretion machinery(45), these results provide an avenue for future research toward elucidating how species may modify alginate production to adapt to diverse environmental niches.

### Genomic Proximity Networks of Acetylated-Cellulose Operons Reveals Duplication of Copolymerase Subunits and Sequence Homology of Loci with Alginate Acetylation Machinery

From the genome sequences surveyed, only four species were identified as possessing acetylated-cellulose operons, comprising two distinct operon clusters with differing operon constitutions among three *Pseudomonas* spp. and *Bordetella avium* 197N (**Supplemental Figure 6**). Contrary to cellulose phylogenetic clusters, the polysaccharide synthase, *wssB*, was divided into distinct Gamma- and Beta-proteobacterial clusters. We also found a distinct phylogenetic cluster identifying a unique tandem duplication of *wssC* in *Bordetella avium* 197N, which was not observed among orthologous cellulose *bcsB* copolymerase loci (**Supplemental Figure 6 (ii)**). This observation might suggest a divergent mechanism of action of cellulose inner-membrane transport. As we previously observed (**Figure 1**), 3 out of 4 of the predicted acetylated-cellulose operons were also found to co-occur with alginate operons. Additional HMM-searches identified significant sequence similarity between acetylated-cellulose *wssBCDE* operon sequences to those previously identified as *bcsABZC*, as well as between acetylated-cellulose acetylation-machinery and their functional homologs in alginate operons (WssH – AlgI; WssI – AlgJ/AlgX). Taken together, these findings suggest that acetylated cellulose production has likely evolved through the duplication and operonic acquisition of the alginate acetylation machinery loci.

### Sequence Variability of Phylogenetic Clusters Reveals Different Degrees of Structural Conservation of Cellulose Biosynthesis Machinery

With the availability of a crystal structure for the BcsA-BcsB inner membrane complex responsible for cellulose biosynthesis and transport(46), we examined the potential structural and functional consequences of the sequence variability of the BcsA and BcsB phylogenetic clusters highlighted above (**Figure 3**). In brief, we generated multiple sequence alignments of eight BcsA and BcsB sequences summarizing the evolutionary diversity of cellulose operon clades identified in **Figure 3**. Residue conservation information from this alignment were subsequently mapped onto the structure of the BcsA-BcsB complex (PDB ID:4HG6 (47); **Supplemental Figure 7**). The results of the following analysis are also consistent when including all predicted BcsA and BcsB sequences. We identified a high degree of sequence conservation among BcsA sequences corresponding to the GT domain responsible for cellulose polymerization. Conserved residues mapped specifically to a cleft in the GT domain where a uridine diphosphate (UDP) carrier moiety is bound and oriented through a conserved QxxRW motif to enable polymerization of glucose monomers of the growing cellulose chain(46). Conversely, the PilZ domain of BcsA, involved in regulation of the GT function in response to c-di-GMP levels shows low conservation overall, except for the subset of residues required for c-di-GMP binding. Further, the periplasmic region of BcsB shows low sequence conservation overall, aside from a number of highly conserved residues in the carbohydrate binding and ferredoxin domains, one of which (L193 of the *Rhodobacter sphaeroides* ATCC 17025 reference sequence) is a putative cellulose binding residue oriented in close proximity to the growing cellulose chain near the exit of the BcsA IM translocation channel. From phylogenetic sequence clustering, structurally relevant conservation features of the cellulose synthase complex can be identified which should facilitate further investigation of cellulose EPS production across phylogenetically diverse species. For example, c-di-GMP binding residues of the PilZ domain of BcsA vary in conservation across phylogenetic clusters, which could impact the binding affinity and limit access of activated glucose monomers to the GT domain, thus limiting the rate of cellulose polymerization. Insertion/deletion events are also observed across BcsB phylogenetic clusters that may facilitate the recruitment of additional periplasmic processing proteins(48), or macromolecular assembly of the BcsA-B complex(49), resulting in differences in the higher-ordered structuring of cellulose microfibres as a consequence of adaptation to diverse environmental niches. These results demonstrate how the application of our phylogenetic clustering methodology can be further extended to provide biologically informative insights into the function of other components of EPS secretion machineries.

### Phylogenetic Clustering Elucidates the Structural and Functional Divergence of the *pgaB* Locus, Revealing the Evolution of PNAG Production Across Gram-negative and Gram-positive Bacteria

PNAG production is found across phylogenetically diverse species and is carried out by the *pgaABCD* operon of Gram-negative(29) and *icaADBC* operon of Gram-positive (50) bacteria. Although the functional and immunological properties of *pga* and *ica* produced PNAG appear to be similar(54), there are important differences between the roles of *pga* and *ica* operon loci(53). Common to both operons is the presence of an integral membrane GT locus, *pgaC* and *icaA*, which are both members of the GT-2 family and share sequence homology(53). In addition, non-homologous loci encoding integral membrane proteins, *pgaD* and *icaD*, are also present and required for the full function of their respective GTs(54,55). In Gram-negatives, PNAG production is regulated through physical interactions between PgaD and PgaC which are stabilized by the allosteric binding of c-di-GMP(56), while in *Staphylococci* PNAG production does not depend on c-di-GMP and is likely regulated by an alternate signaling pathway(57). Deacetylation of PNAG is carried out by *pgaB* and *icaB* loci and has been shown to play a crucial role in biofilm formation and immune evasion(52,58). *pgaB* also possesses an additional C-terminal glycoside hydrolase domain which cleaves the PNAG polymer following its partial deacetylation(59), although the mechanism of how these activities are coordinated and the biological role of the hydrolase activity is unknown. Unique to *pga* operons is a loci encoding an outer membrane export pore, *pgaA(60)*, and in *ica* operons an additional integral membrane protein, *icaC*, which has been proposed to be involved in PNAG O-succinylation(53). Using Gram-negative *pga* loci as seed sequences for the reconstruction of synthase-dependent PNAG operons, we were also able to identify Gram-positive *ica* operons based on significant sequence similarities to *pgaB* and *pgaC* loci. Our phylogenetic clustering approach also revealed that *pgaC/icaA* sequences clustered into a single clade, while *pgaB/icaB* were associated with distinct sequence clades (**Supplemental Figure 4**). To explore the evolution of Gram-negative and Gram-positive *pga* and *ica* operons, we generated multiple sequence alignments for representative sequences of 18 PgaB clades. Our phylogenetic clustering results confirm previous observations(53) that the glycoside hydrolase domain is exclusively associated with Gram-negative *pga* operons (PgaB_G1) and is absent in a clade of *Staphylococcus* Gram-positive *ica* sequences (PgaB_G3) (**Supplemental Figure 8A**). We also identified additional Gram-positive *icaB* clades among non-*Staphylococcus* spp., e.g. *Bacillus, Lactococcus*, and *Mycobacterium* (**Supplemental Table 1**), which possess operons lacking the *icaC* locus(53). Interestingly, we also identified a number of divergent Gram-negative *pgaB* clades resembling *icaB* clade sequences. Members of these clades lacked the canonical N-terminal glycosyl hydrolase domain, and were distinguished by possessing N-terminal fusions, primarily of GT domains. Furthermore these *pgaB* clades are associated with operons lacking detectable *pgaA* outer membrane pore locus and *pgaD* (**Supplemental Figure 4 (v)**). Although PNAG production in these species has not been experimentally confirmed, these findings suggest if the polymer is produced it is under a novel mode of regulation by c-di-GMP, that glycoside hydrolase activity might not be essential for PNAG export across all Gram-negative species, and that other modes of export may exist. The fusion of would also suggest that the de-acetylase activity of PgaB in these organisms may be associated with the periplasmic face of the inner membrane, in contrast to dual domain PgaB clades where the protein is predicted to function at the periplasmic face of the outer membrane(60).

In addition to these novel domain fusion events, PgaB phylogenetic clustering enabled us to resolve distinct events affecting the evolution of the deacetylase domain across different operon clades. Using the *E. coli* K12 MG1655 sequence of the largest PgaB clade (PgaB_G1) as a reference, multiple sequence alignments against other representative PgaB clade sequences identified several regions of insertion/deletion events (**Supplemental Figure 8A**). When these regions were mapped to the published crystal structure of PgaB (PDB ID: 4F9D(61)), they were found to correspond to distinct structural elements surrounding the conserved deacetylase core (**Supplemental Figure 8B-C**). We assigned insertion/deletion regions a number according to their order of appearance in the multiple sequence alignment of PgaB deacetylase domains, and divided them into two categories (**Supplemental Figure 8D**). The first two indel regions, 1 and 2, resided in the N-terminal region of the reference *E. coli* sequence, and corresponded to beta-strands flanking the conserved active site residues involved in deacetylation, His55, Asp114, and Asp115. Region 1 was associated with Gram-positive *icaB* and comprised insertions of ∼10aa in *Staphylococcus aureus* VC40 (PgaB_G3), as well as *Bacillus infantis* NRL B-14911 (PgaB_G7), *Lactobacillus plantarum* 16 (PgaB_G9), *Leptospirillum ferriphilum* ML-04 (PgaB_11). Structural characterization of *Ammonifex degensii* IcaB (PgaB_G3) identified residues overlapping with Region 1 as encoding a hydrophobic loop responsible for membrane localization in this species(62). Region 2 was found to be exclusive to Gram-negative *pgaB* loci and comprised a much larger insert of ∼77aa in *Geobacter metallireducens* GS-15 (PgaB_G2), *Crinalium epipsammum* PCC 9333 (PgaB_G5), and *Colwellia psychrerythraea* 34H (PgaB_6). The functional role of this insert is unknown.

The last three insertion/deletion regions, 3-5, occurred in a region oriented away from the deacetylase active site, and correspond to two beta-turn motifs and an alpha-helix cap, respectively. To further elucidate the biological import of identified PgaB indel regions, we examined regions 3, and 5 in the context of Gram-negative PNAG modification. In the *E. coli* K12 MG1655 PgaB_G1 sequence, region 3 encompasses a beta-turn with an elongated loop, which is spatially proximal to a disordered loop and alpha helix (pos. 367-392) on the N-terminal region of the PgaB glycoside hydrolase domain. Region 3 also encodes a histidine (*E. coli* PgaB - H189) which is part of the Ni binding pocket of Gram-negative PgaB deacetylases. Both regions contain polar and electrostatically charged residues which are highly conserved across PgaB_G1 sequences (**Supplemental Figure 8E**). Region 5 corresponds to an 8 amino acid elongation of an alpha-helix (pos. 219-226), which also appears to provide an additional point of contact between the deacetylase and hydrolase domains. Although region 5 is also shared with *icaB* associated sequences (PgaB_G3), region 3 appears only in other dual deacetylase-hydrolase Gram-positive *pgaB* sequences identified in the sporulating bacteria *Lachnoclostridium phytofermentans* ISDg and *Kitasatospora setae* KM-5043. Although initial PFAM searches failed to identify the additional Gram-positive C-terminal domains, subsequent BLAST searches revealed them to be homologous to glycoside hydrolases. In region 4 a unique 29 amino acid insertion was also identified in *Lachnoclostridium phytofermentans* ISDg (PgaB_G16), which may play a compensatory role for the absence of 9aa in region 3. These insertion regions suggest an overall functional importance in ensuring stability between each domains and could play a role in coordinating their activities. These findings in combination with our identification of *ica-*like operon organizations among environmental Gram-negative species (**Supplemental Figure 4(v)**) suggest that Gram-negative *pga* operons may share a common evolutionary origin with Gram-positive *ica* operons. Recent research is providing growing evidence for the emergence of the di-derm Gram-negative architecture from sporulating monodermal Gram-positives (64), which provides a plausible evolutionary context for the insertion/deletion events observed among *pgaB/icaB* deacetylase domains. Through the loss of inner membrane localization(62) (Region 1), the compensatory gain of an N-terminal palmitoylation site(54), along with a C-terminal fusion of a hydrolase domain (Regions 3-5), an ancestral deacetylase locus may have been adapted to regulate the export of PNAG(54) at the outer membrane of Gram-negative *pga* operon lineage.

## DISCUSSION

In this work we describe a novel and generalizable approach for the systematic classification and presentation of bacterial protein families in the context of their host operon. Protein families are defined as sets of homologs (groups of related sequences having a common evolutionary ancestor) sharing a particular set of sequence motifs or structural domains that can be utilized to determine their biological roles. For example, the PFAM database utilizes curated sets of protein family sequences in the generation of profile HMMs(64). A key challenge that complicates the definition of these relationships are evolutionary events such as duplication, gene fusion, and HGT. In attempts to account for such events, a variety of computational approaches have been developed for refining functional assignments either by graphical clustering of pair-wise protein sequence similarities (e,g, COG(65), OrthoMCL(13) and EggNOG(14)), or through the generation of hierarchical evolutionary relationships and construction of phylogenetic trees (e.g. TreeFAM(66) and TreeCL(67)). However, these methods are limited in their ability to provide further resolution of sequence diversity within a family that might otherwise offer additional insights into evolutionary events that allow taxa to adapt to specific environments.

Agnostic approaches to define sub-clusters of evolutionarily related protein families have ranged from phylogenetic tree reconstructions (68) to hierarchical clustering of pairwise global sequence alignments(69). Here we present an extension of previous efforts, and introduce a novel systematic approach for defining protein sub-family relationships through the clustering of phylogenetic trees. Key to this approach is defining a scoring function that allows a phylogenetic tree to be resolved into optimal clusters that best capture the similarities between cluster members, as well as the dissimilarities between clusters. Combining two clustering quality metrics (Silhouette and Dunn index) and proportion of sequences clustered, we demonstrate that our approach is able to classify a diverse array of operon-associated protein families into taxonomically consistent and functionally informative sub-clusters. Genomic-proximity networks were also constructed to provide an intuitive means of utilizing phylogenetic clusters to examine diverse mechanisms of operon evolution across taxonomically diverse bacterial genomes. Genomic-proximity networks have previously been utilized for inferring functional relationships(70), understanding mechanisms underlying bacterial genomic organization into functionally related gene clusters(71), and transcriptional regulation of bacterial operons(72). In this study we extend the application of genomic-proximity networks as a tool for the systematic exploration of operon evolution resulting from locus divergence, loss, duplication, and rearrangement events.

To demonstrate the effectiveness of our approach, we applied our methods to classify the stynthase-dependent bacterial EPS operon machineries for 5 different polymers: cellulose, acetylated-cellulose, alginate, Pel and PNAG. There has been only one previous attempt to classify synthase dependent EPS operons and this focused specifically on the cellulose system(25). In that study, cellulose operons were categorized into four major types, based on the presence or absence of experimentally validated accessory loci involved in cellulose production. Here, we based our analysis on the four core operon loci, *bcsABZC*, deemed essential for cellulose production. Cellulose operon clades identified in this study showed little consistency with the previously defined four major cellulose operon types(25), suggesting that the conservation of accessory loci is more variable across bacterial species compared to loci encoding core EPS functionalities. However, one operon type was identified in this analysis, representing the loss of the BcsC outer membrane transporter identified among a subset of alpha-proteobacterial genomes, which include several known cellulose producing species(47,73) suggesting a novel mechanism of cellulose export(**Figure 3(iii)**)(25). We also found that the loss of BcsC has resulted in an increased divergence of BcsB loci in these genomes, which highlights the key role of BcsB as an intermediary between cellulose biogenesis and periplasmic transport (**Figure 6**).

In general, inner membrane components involved in EPS polymerization were found to be relatively conserved across all systems examined, while periplasmic and outer-membrane components showed a relatively increased degree of evolution, which are likely to have important functional implications. For example, in the cellulose and Pel operon networks (**Figures 3 and 5, and Supplemental Table 4**), rearrangement events involving the periplasmic glycosyl hydrolase (BcsZ) and glycosyl hydrolase/deacetylase (PelA) were found to be a defining feature of several operon clades. It is interesting to note that these rearrangements have resulted in a change in the ordering of *bcsZ* and *pelA* relative to their respective outer-membrane transport pore loci, which highlights the important role of polysaccharide modification in both the biogenesis and regulating extracelluar EPS transport(20,38,74). Similarly, the rearrangement of alginate modification machinery loci (*algIJF*) was observed as a distinguishing feature of *Pseudomonas* spp. operon clades. These findings suggest that rearrangement and locus ordering may serve as an important means of regulating EPS production by modifying the timing of translation of modification enzymes, which could affect the assembly of EPS complexes or the structural properties of EPS produced (37,49,75).

Furthermore, identifying operon clades through a phylogenetic approach elucidated numerous instances of cellulose whole operon duplications arising from HGT of two evolutionary distinct operon clades (**Figure 4**). Such large-scale duplications, if they are functional, may either serve as a dosage response to given environmental stressors, as observed in the duplication of bacterial multiple-drug transporter operons(76), or could be under the regulation of different environmental stimuli. Interestingly, representative species of the two cellulose operon lineages identified in HGT events, e.g. the plant and human pathogens, *D. dadantii* and *S. enterica*, respectively, are known to produce structurally distinct forms of cellulose with different properties and roles in pathogenesis(77,78). Furthermore, we identified that BcsB divergence was also seen to accompany the rearrangement or horizontal transfer of these operons, which further suggests that it may play a key role in the fine-tuning of cellulose production by coordinating the export of growing cellulose polymers through the periplasm. Furthermore, our analyses of acetylated-cellulose, alginate and PNAG operons suggest a dynamic evolutionary scenario for the evolution of EPS biofilm production through the acquisition of novel polysaccharide modification loci. The limited number of acetylated-cellulose operons identified, their frequent co-occurence in alginate possessing species, and significant sequence similarities between acetylation machinery loci, suggests that the cellulose acetylation machinery is likely to have originated from previously existing alginate operons in *Pseudomonas* spp. The evolutionary trajectories of Gram-positive and Gram-negative PNAG operon lineages appears to have resulted through the fusion of glycosyl hydrolase and deacetylase domains in Gram-negative *pgaB* loci..

A further key finding from this study was the identification of homologous *pel* operons in the genomes of several Gram-positive bacteria. With the additional identification of homologs of PelD through iterative HMM searches, our analyses have uncovered a novel example of c-di-GMP regulation of biofilm machinery in Gram-positive bacteria. In the accompanying paper we experimentally validate that a predicted *pel*-like operon in *B. cereus* ATCC 10987 is responsible for biofilm production which is regulated bythe binding of c-di-GMP to PelD (Whitfield et al submitted).

Together this work demonstrates a novel integrative approach combining phylogenomics and genomic-context approaches to systematically explore the adaptive implications of sequence divergence of protein families associated with operon associated EPS secretion machineries. Further extension of this work holds great potential as a general approach for elucidating how bacterial operon encoded biological pathways and complexes have contributed to bacterial adaptation to and survival in diverse environmental niches and lifestyles.

## METHODS

### Sources of Data

Sequences corresponding to experimentally characterized EPS operon loci were obtained from the National Centre for Biotechnology Information (NCBI) reference sequence database(79) (**Supplemental Table 3**). Fully sequenced genomes and associated protein sequences were obtained for 1861 bacteria from the NCBI (Retrieved April 20th 2015) (**Supplemental Table 6**). For each bacterial strain predicted to possess an EPS operon, metadata corresponding to niche (host-associated or environmental) and lifestyle (pathogenic or non-pathogenic) were collated from literature searches (**Supplemental Table 7**).

### Prediction of EPS operons

To identify putative EPS operons, we applied an iterative HMM-based sequence similarity profiling strategy. For each set of EPS loci, we first constructed a HMM; alignments were constructed using MUSCLE v.3.8.1551(80), with default settings, from which HMM-models were built using HMMER v.3.1b2(81), with default settings. Each HMM was then used to identify additional EPS loci within the set of 1861 bacterial genomes. The 20 non-redundant sequences (as defined by >97% sequence similarity; i.e. to eliminate sequences from closely related strains) that had the highest scoring matches (as defined by e-values) were then retrieved and added to the original set of loci to construct a new set of HMMs. Using these new sets of HMMs, sets of EPS loci for the reconstruction of EPS operons (see below) were predicted through sequence similarity searches of the 1861 genomes using HMMER, with default settings. Significant sequence matches were defined as those with E-values <= 1e-5.

To reconstruct putative EPS operons from the sets of loci retrieved from our searches, we first retrieved locus start and stop positions for each locus from their RefSeq entry. We then define putative operons using the following two rules: first only loci that occur within a distance of twice the size of a reference EPS operon to other loci are considered; second intergenic distances of individual loci must be <= 5 Kbp; third putative operons must consist of at least one locus encoding a putative polysaccharide synthase, together with at least one other locus. To detect previously undiscovered loci that may have been missed in the first rounds of HMM searches, predicted loci of reconstructed operons were used to generate expanded locus-specific HMM models and were subjected to an additional round of HMM searches. This process was performed using custom Perl scripts and results in a list of predicted EPS operons identified in each of the 1861 genomes.

### Classification of Evolutionary Events

For each EPS system (cellulose, acetylated-cellulose, PNAG, pel, and alginate), the locus assignments of each reconstructed operon was compared to a defined reference EPS operon compositions and locus ordering (**Supplemental Table 4**) and were classified into the following evolutionary events; 1) locus losses - the total number of reference loci missing or not detected by HMM searches; 2) locus duplications – number of distinct loci appearing as multiple significant hits to the same HMM model < 10kBP apart; 3) locus fusions – the number of loci that were significant hits to two or more reference EPS locus HMM models; 4) operon rearrangements – the number of predicted operons with locus ordering (accounting for transcriptional direction) different from the reference operon; 5) operon duplications – number of predicted operons (as defined above) present in the same genome >= 10 Kbp apart.

### Classification of EPS loci

Systematic classification of each EPS operon family starts with first merging closely related sequences using CD-HIT v.4.6.3(82) with default settings (using global sequence identity threshold 0.9; word length 5) to generate a non-redundant set of sequences for each family. Multiple sequence alignments (MSAs) were then generated using MUSCLE and trimmed using trimal v.1.2rev59(83) (using - automated1 setting). The resulting alignment was then used to construct a phylogenetic tree using PhyML v.3(84), with default parameters (LG substitution model, with 1000 bootstrap replicates). For each tree, an optimal set of clusters is then generated by traversing the tree, starting at the tips and iteratively increasing evolutionary distance (defined as the number of expected amino-acid substitutions per site) between branches. At each step an evolutionary distance cutoff threshold is chosen (beginning from 0 to the maximum distance for a given tree and increasing in increments of 0.01) and all sequences which share a branch less than the given threshold are assigned to the same evolutionary cluster. This results in the generation of increasingly coarse clusters of sequences with increasing sequence dissimilarity, such that in the final step all sequences are assigned to a single cluster. At this stage, for all possible clusterings three metrics are calculated and summed together to calculate a clustering quality score: (1) proportion of sequences clustered (p) number of sequences clustered / total number of sequences); (2) the average silhouette score (s_avg) (85):

For each sequence, i, its silhouette score, s(i), is defined as: 

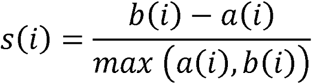

Where a(i)=average evolutionary distance (expected number of substitutions per site) i) is the lowest average evolutionary distance to any other cluster of which i is not a member; and (3) Dunn index (DI)(86), for a set of m clusters, its Dunn index, DI, is defined as: 

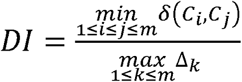

Where DI is the evolutionary distance between clusters i and j and Δc is the size of cluster c. Note that a higher s(i) indicates that a sequence is well matched to other members of its cluster and not well matched to neighbouring clusters. Furthermore, a higher DI indicates clusters that are compact (smaller cluster sizes) and well differentiated (larger inter-cluster distances). Thus, the evolutionary distance cutoff which maximizes p + s_avg + DI is chosen as the optimal phylogenetic clustering for a given set of EPS locus sequences.

### Construction of EPS Operon Genomic-Proximity Networks

To visualize evolutionary and genomic organization relationships of predicted EPS operons, genomic proximity networks were generated in which each node represents an individual EPS locus cluster (as defined above), and an edge connecting a pair of nodes represents the average genomic distance (base pairs) between loci represented by each node found in the same genome. Further, nodes are represented as pie-charts indicating phylogenetic distribution of each EPS locus, as defined by NCBI taxonomic classification scheme. Networks were visualized using Cytoscape (version 3.5)(87).

## ACKNOWLEDGEMENTS

JP and CB-T were supported by grants from the Natural Sciences and Engineering Research Council (RGPIN-2014-06664 & RGPIN-2019-06852) and the National Institutes of Health (R21AI126466). This work was also supported in part by grants from the Canadian Institutes of Health Research (CIHR) (MOP 43998 and FDN154327 to PLH). PLH is a recipient of a Canada Research Chair. GBW and LSM have been supported by graduate scholarships from the Natural Sciences and Engineering Council of Canada (NSERC). GBW has been supported by a graduate scholarship from Cystic Fibrosis Canada. LSM has been supported by graduate scholarships from the Ontario Graduate Scholarship Program, and The Hospital for Sick Children Foundation Student Scholarship Program. Computing resources were provided by the SciNet HPC Consortium; SciNet is funded by: the Canada Foundation for Innovation under the auspices of Compute Canada; the Government of Ontario; Ontario Research Fund–Research Excellence and the University of Toronto.

## FIGURE LEGENDS

**Supplemental Figure 1. Lifestyle and Niche Distribution of Predicted EPS Operons**. The number of bacterial genomes with different combinations of predicted EPS operons, further represented with their distribution (% bacterial genomes) across different lifestyles and environmental niches. Asterisks indicate statistically significant enrichment of single or multiple EPS operon combinations among pathogenic (red asterisks) or non-pathogenic bacteria (green asterisks) (one sided T-test p <= 0.05 with Bonferroni correction).

**Supplemental Figure 2. Species Diversity of Predicted Synthase Dependent EPS Systems (Shannon Diversity)**.

**Supplemental Figure 3. Identification of Gram-positive *pel* Operons**. (A) Subnetwork depicting Gram-positive *pel* operon clades with varying numbers of loci identified as significant matches (e-value < 1e-5) in first-pass (unfilled nodes) and iterative HMM searches (grey nodes). Selected examples shown: (i) PelA-PelFG sequences identified by first-pass HMM hits; (i.b) Iterative HMM searches identifying additional *pelA* loci in *B. cereus* ATCC 10987, a known pellicle producing Gram-positive; (ii) Additional *pelD* loci identified by iterative HMM; (iii) Gram-positive pel operons with only *pelF* and *pelG* loci identified. (B) Operon organizations of selected examples of Gram-positive *pel* operons (corresponding highlighted in panel A) with additional highly divergent loci identified (red boxes: hits above HMM e-value threshold of 1e-5).

**Supplemental Figure 4. Genomic-Proximity Network of Phylogenetically Clustered *pga* Operons**. Phylogenetically clustered operon loci are arranged according to the canonical *pga* operon ordering indicated by the grey sidebar. Inset boxes depict selected examples of *pga* operon clades distinguished by evolutionary events: i) Divergence of *pgaD* corresponding to related enterobacterial species including pathogen-specific losses of *pgaA* and *pgaB* loci critical for PNAG export; ii) Operon duplications occurring in aquatic niche dwelling bacteria, including a partial duplication of the *pga* operon specific to the opportunistic pathogen *Acinetobacter baumannii* spp. and a whole operon duplication identified in *Methylovora versatilis*; iii) A unique *pga* operon organization among environmental bacteria lacking a *pgaD* locus; iv) Gram-positive *ica* operons (annotated by their HMM hits to corresponding Gram-negative *pga* loci) with divergent *icaB* loci, resulting from novel domain acquisitions (iv.b and iv.c); v) A novel *pga* derived operon resulting from multiple tandem duplications of the *pgaC* polysaccharide synthase and lack of detectable *pgaA* outer membrane pore and *pgaD*. Node size indicates the relative number of sequences per phylogenetic cluster; node colouring represents the taxonomic distribution of loci for a given cluster; edges connect clusters which co-occur in the same genome(s); edge colour indicates the genomic-proximity of loci clusters. The network was visualized using Cytoscape 3.5.1(87).

**Supplemental Figure 5. Genomic-Proximity Network of Phylogenetically Clustered Alginate Operons**. Phylogenetically clustered operon loci are arranged according to the canonical alginate operon ordering indicated by the grey sidebar. Inset boxes depict selected examples of alginate operon clades distinguished by evolutionary events: Inset boxes depict selected examples of alginate operon clades distinguished by evolutionary events: i) Canonical alginate operon organization with a partial operon duplication event identified in *Pseudomonas resinovorans* 136 resulting in the loss of alginate acetylation machinery (ib – indicated by A*); ii) A distinct alginate operon clade (ii.a-c) identified by rearrangement of acetylation machinery (indicated by B*) as well as HGT events with canonical alginate operon possessing species; iii) Atypical alginate operons involving loss of outer membrane transport loci or portions of acetylation machinery in deep sea dwelling bacteria. Node size indicates the relative number of sequences per phylogenetic cluster; node colouring represents the taxonomic distribution of loci for a given cluster; edges connect clusters which co-occur in the same genome(s); edge colour indicates the genomic-proximity of loci clusters. The network was visualized using Cytoscape 3.5.1(87).

**Supplemental Figure 6. Genomic-Proximity Network of Phylogenetically Clustered Acetylated-Cellulose Operons**. Phylogenetically clustered operon loci are arranged according to the canonical acetylated cellulose operon ordering indicated by the grey sidebar. Inset panels identify three acetylated-cellulose operons identified in *Pseudomonas* spp. (i) and a single *Bordetella avium* genome possessing a duplicated polysaccharide co-polymerase *wssC* locus (ii - indicated by red asterisk). Node size indicates the relative number of sequences per phylogenetic cluster; node colouring represents the taxonomic distribution of loci for a given cluster; edges connect clusters which co-occur in the same genome(s); edge colour indicates the genomic-proximity of loci clusters. The network was visualized using Cytoscape 3.5.1(87).

**Supplemental Figure 7. Phylogenetic Sequence Clustering Reflect Differences in Structural Conservation Between Cellulose Synthase Complex Subunits BcsA and BcsB**. Top panel - Sequence conservation was mapped onto the cellulose synthase complex, BcsA-BcsB (4HG6 – *Rhodobacter sphaeroides* ATCC 17025) comprising sequences from eight species representing distinct cellulose operon clades (**Figure 4(i)-(iv)**). Lower panels - structural and multiple sequence alignments indicate a high degree of conservation corresponding to BcsA glycosyl hydrolase catalytic core domain and regions of the cellulose translocation channel (i) and UDP binding sites of the BcsA PilZ domain (ii). In Contrast, low overall sequence conservation is found among the carbohydrate binding and ferredoxin domains (CBD1-2, and FD1-2) of BcsB sequences, except the highly conserved cellulose binding site residing in CBD-2 (iii). The translocated cellulose polymer is indicated in green. BcsA domains identified using PFAM predictions for the *R. sphaeroides* reference sequence, BcsB domains were assigned according to (45). Multiple sequence alignment was visualized generated using Geneious 10.2.2 (http://www.geneious.com), protein structure was visualized using Chimera 1.11.2(89).

**Supplemental Figure 8. Phylogenetic Clustering Reveals Structural Evolution of PNAG PgaB Periplasmic Modifying Enzyme Distinguishing Gram-Negative and Gram-Postive PNAG Operon Clades**. A) - Multiple sequence alignment of representative sequences comprising all PgaB phylogenetic clusters. Global sequence conservation compared against *E. coli* MG1655 K12 PgaB, phylogenetic cluster PgaB_G1, indicates presence of polysaccharide deacetylase domain (blue box) but an absence of glycosyl-hydrolase domain in non-PgaB_G1 sequences. Red arrows indicate phylogenetic group specific N-terminal domain fusions predicted by PFAM searches; C-terminal domain fusions identified (red box) as putative hydrolase domains from BLAST searches. B) - A close up view of sequence conservation of PgaB polysaccharide deacetylase domains with indel events highlighted: green boxes indicate insertions identified in non PgaB_G1 sequences; teal boxes indicate insertions in PgaB_G1 sequence residing in the C-terminal alpha-helix cap (yellow box). C – Crystal structure of *E. coli* PgaB (4F9D) indicating conservation of the deacetylase domain catalytic core. D – Deacetylase domain with indel regions indicated according to the colour scheme described for panel B. E – C-terminal alpha helical cap region of the PgaB deacetylase domain indicating insertions of the PgaB_G1 region that are spatially proximal to an N-terminal region of the hydrolase domain (light purple); comparison of the same regions with PgaB_G1 sequence conservation indicated. Multiple sequence alignment was visualized generated using Geneious 10.2.2 (http://www.geneious.com), protein structure was visualized using Chimera 1.11.2(89).

