## Supplemental Figure 1 for "A systematic pipeline for classifying bacterial operons reveals the evolutionary landscape of biofilm machineries"

| | | | | | | | | | | | | | |
|---|---|---|---|---|---|---|---|---|---|---|---|---|---|
| Acetylated-Cellulose | + | + | | | | | | | + | | | | |
| Alginate | + | + | + | + | + | | | | | + | | | |
| Cellulose | | | | | | + | + | | | | + | | |
| Pel | | | + | | + | + | | + | | | | + | |
| PNAG | + | | | + | + | | + | + | | | | | + |
| Total Species Genomes | **2** | **1** | **20** | **5** | **2** | **10** | **83** | **7** | **1** | **30** | **274** | **101** | **189** |
| **T-Test Significance Pathogen** vs. **Non-Pathogen** | | | * | | | * | * | | | * | * | * | * |
| Lifestyle: Non-Pathogen | 100 | | 25 | 100 | 100 | 70 | 33 | 86 | | 83 | 56 | 63 | 43 |
| Lifestyle: Pathogen | | 100 | 75 | | | 20 | 66 | | 100 | 17 | 38 | 19 | 56 |
| Lifestyle: Unknown | | | | | | 10 | 1 | 14 | | | 6 | 18 | 1 |
| Niche: Host-Human | | | 55 | | | | 55 | | | 7 | 30 | 16 | 45 |
| Niche: Host-Other | | | 5 | | | | 4 | | 100 | 7 | 6 | 6 | 9 |
| Niche: Microbiome | | | | | | | 1 | | | 3 | 5 | 1 | 2 |
| Niche: Plant | 50 | 100 | | 20 | | 20 | 12 | | | 10 | 11 | 9 | 6 |
| Niche: Rhizosphere | | | 5 | 60 | 100 | 10 | 5 | | | 10 | 11 | 7 | 3 |
| Niche: Food | | | | | | | | | | | 3 | 2 | 3 |
| Niche: Freshwater | | | | | | | 2 | | | 10 | 3 | 2 | 2 |
| Niche: Hot spring | | | | | | 10 | 1 | | | | 1 | 4 | 3 |
| Niche: Industrial | 50 | | 5 | | | | 1 | 14 | | 10 | 5 | 10 | 4 |
| Niche: Lab derived | | | 25 | | | | 13 | | | 10 | 2 | 2 | 5 |
| Niche: Marine sediment | | | 5 | | | | | 29 | | 7 | 3 | 9 | 6 |
| Niche: Seawater | | | | | | 20 | 1 | | | | 5 | 3 | 1 |
| Niche: Soil | | | | 20 | | 30 | 2 | 43 | | 27 | 8 | 12 | 11 |
| Niche: Unknown | | | | | | 10 | 1 | 14 | | | 6 | 18 | 1 |

Host-Associated

Environmental /Other

% of Species Found (Percentile)

-100%

-10%
