## Supplementary figures and images for "A systematic pipeline for classifying bacterial operons reveals the evolutionary landscape of biofilm machineries"

### Supplemental Figure 2

**Phylogenetic Diversity of Predicted EPS Operons**

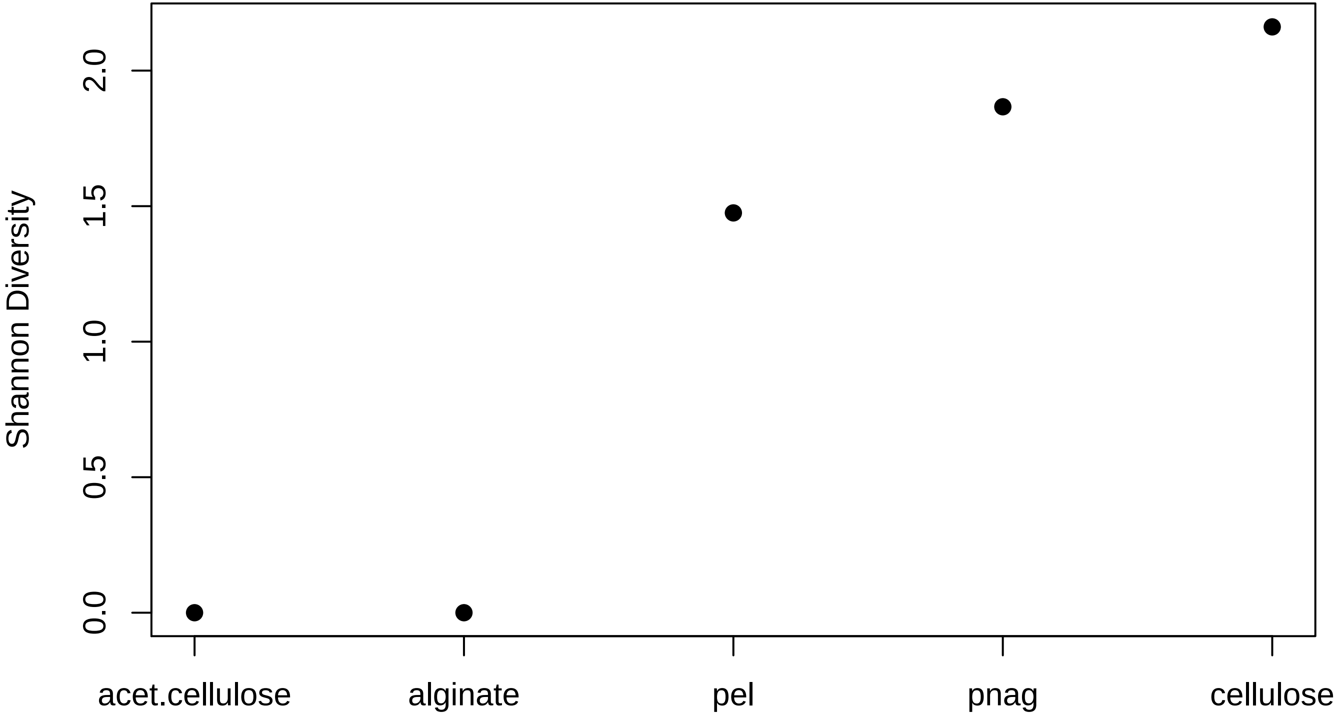

### Supplemental Figure 3

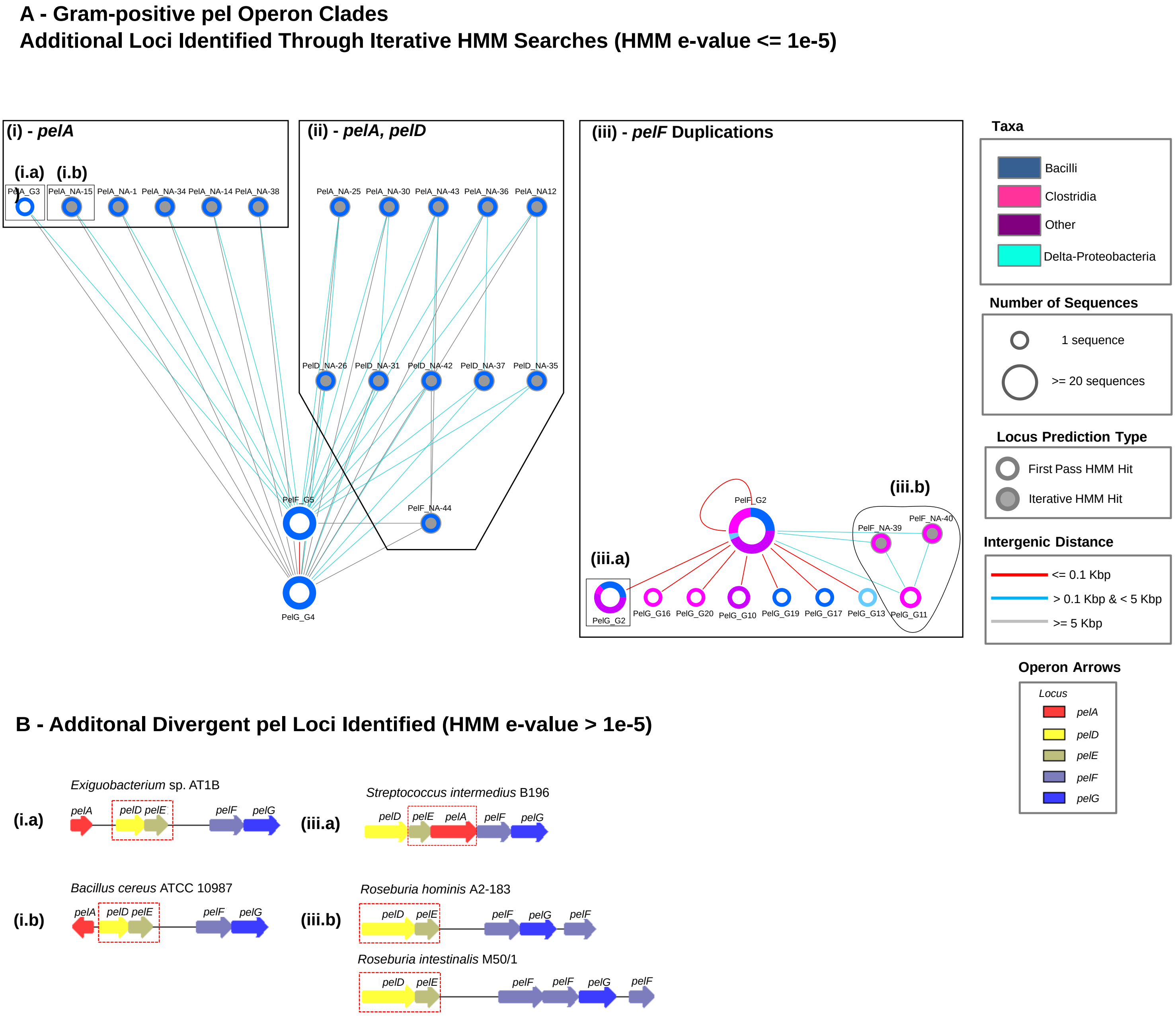

### Supplemental Figure 4

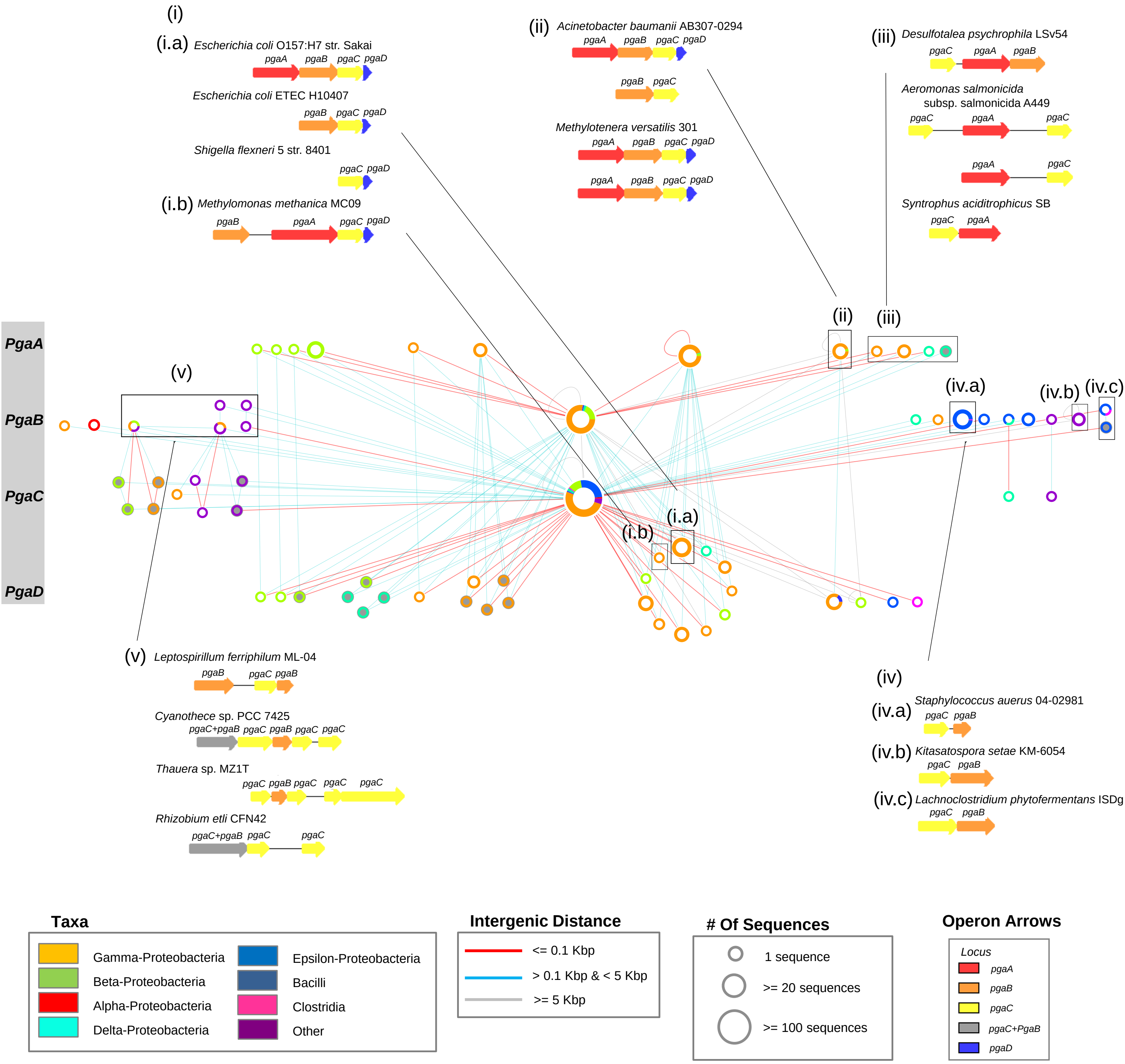

### Supplemental Figure 5

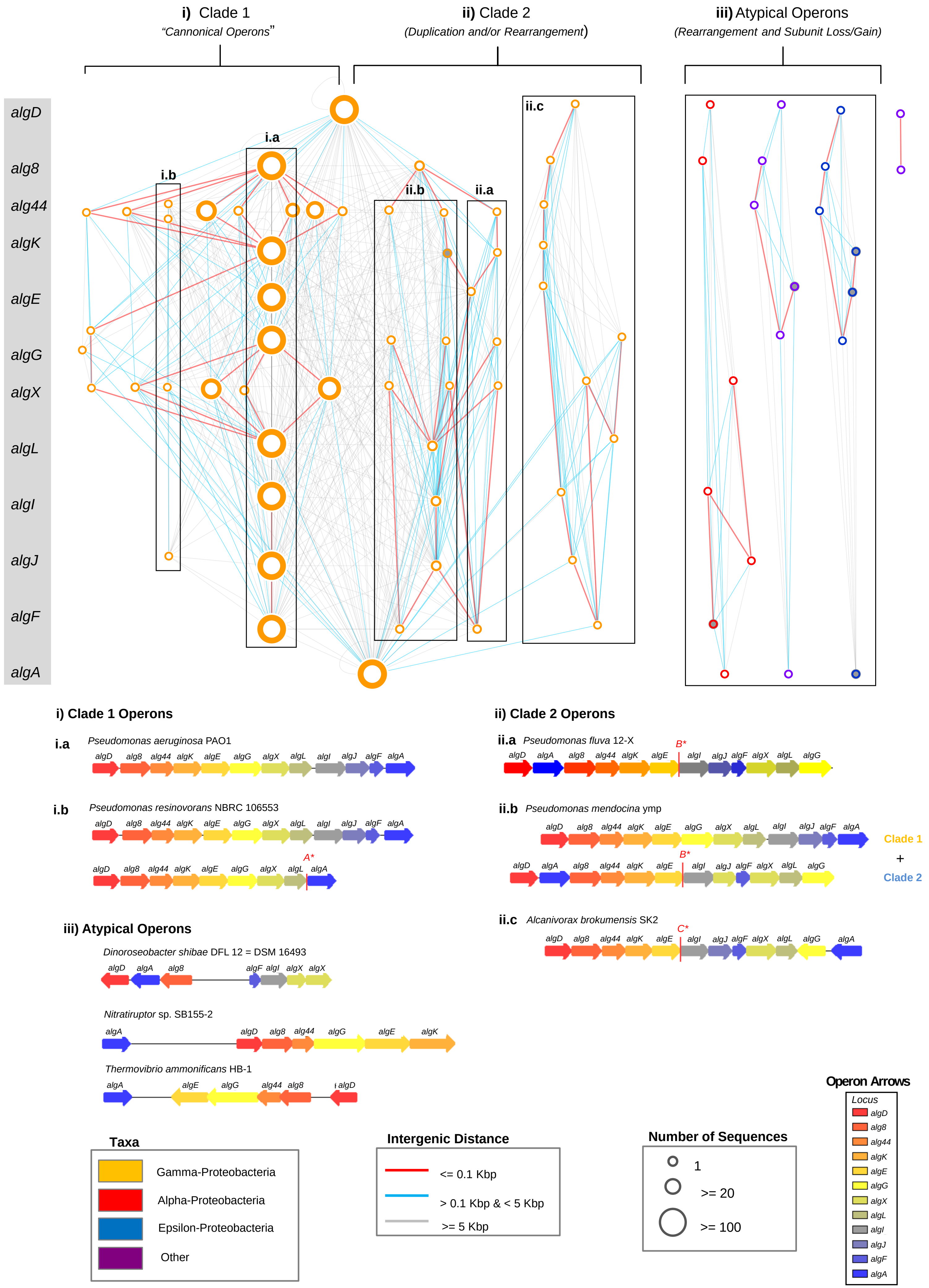

### Supplemental Figure 6

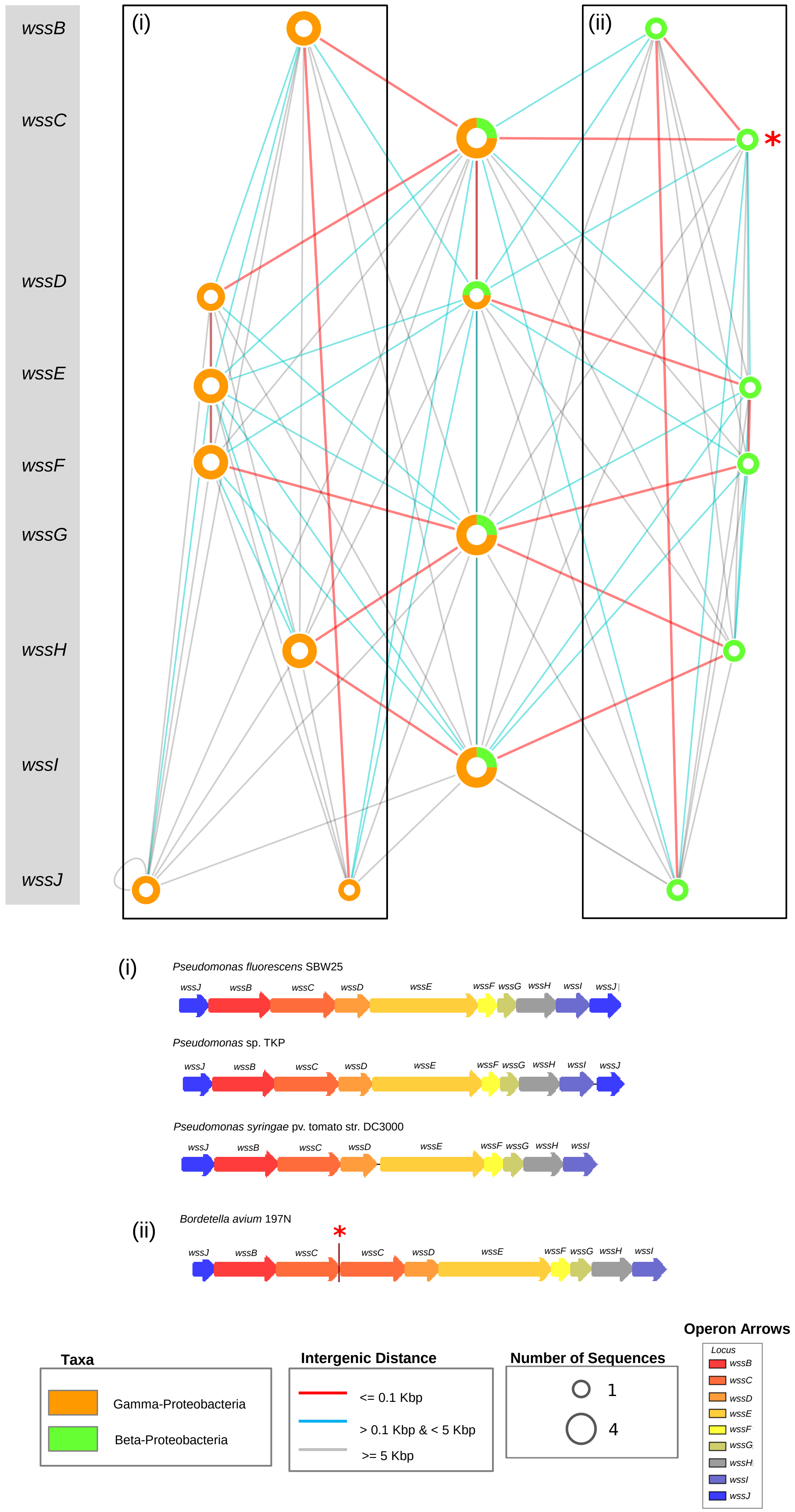

### Supplemental Figure 7

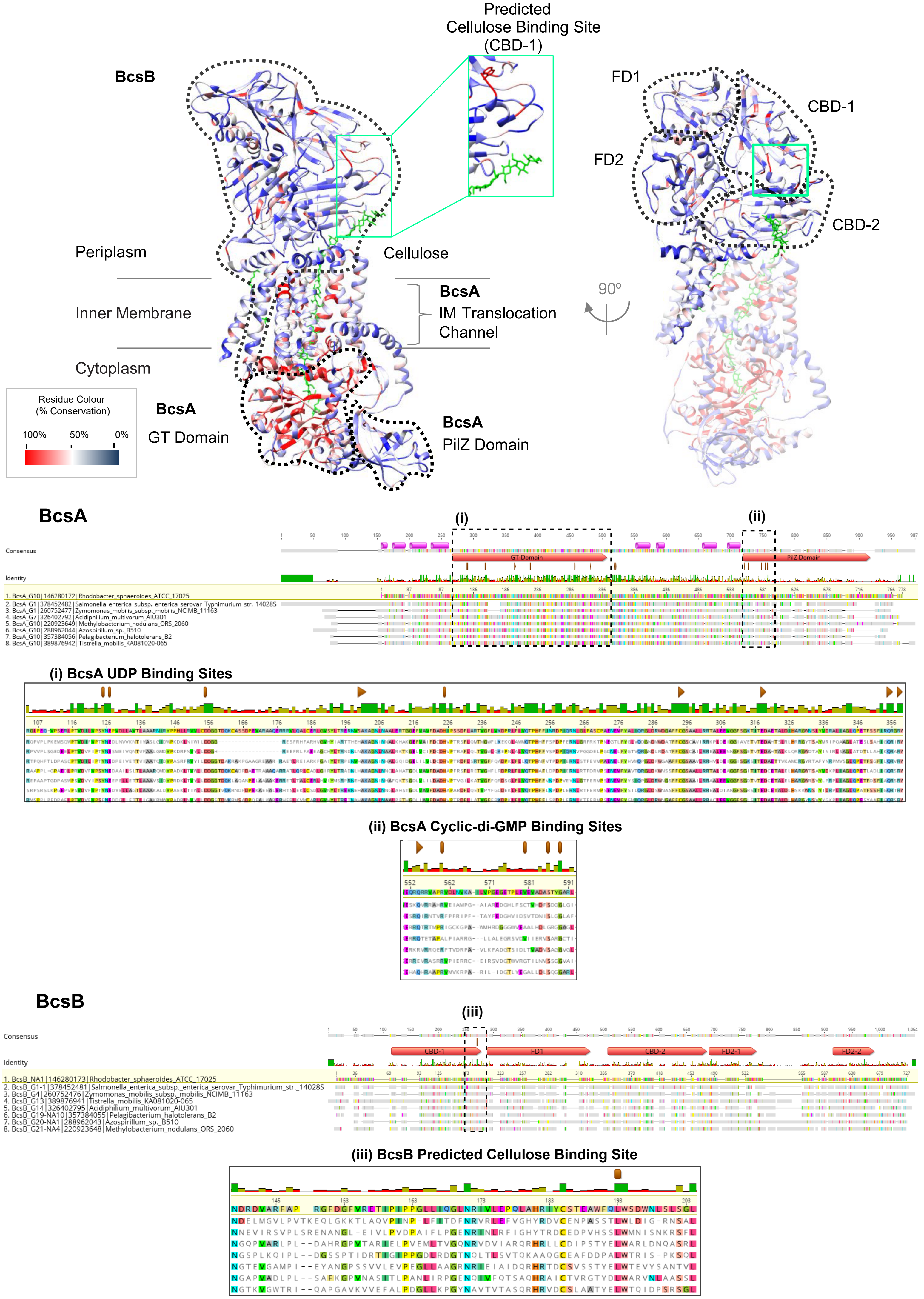

### Supplemental Figure 8

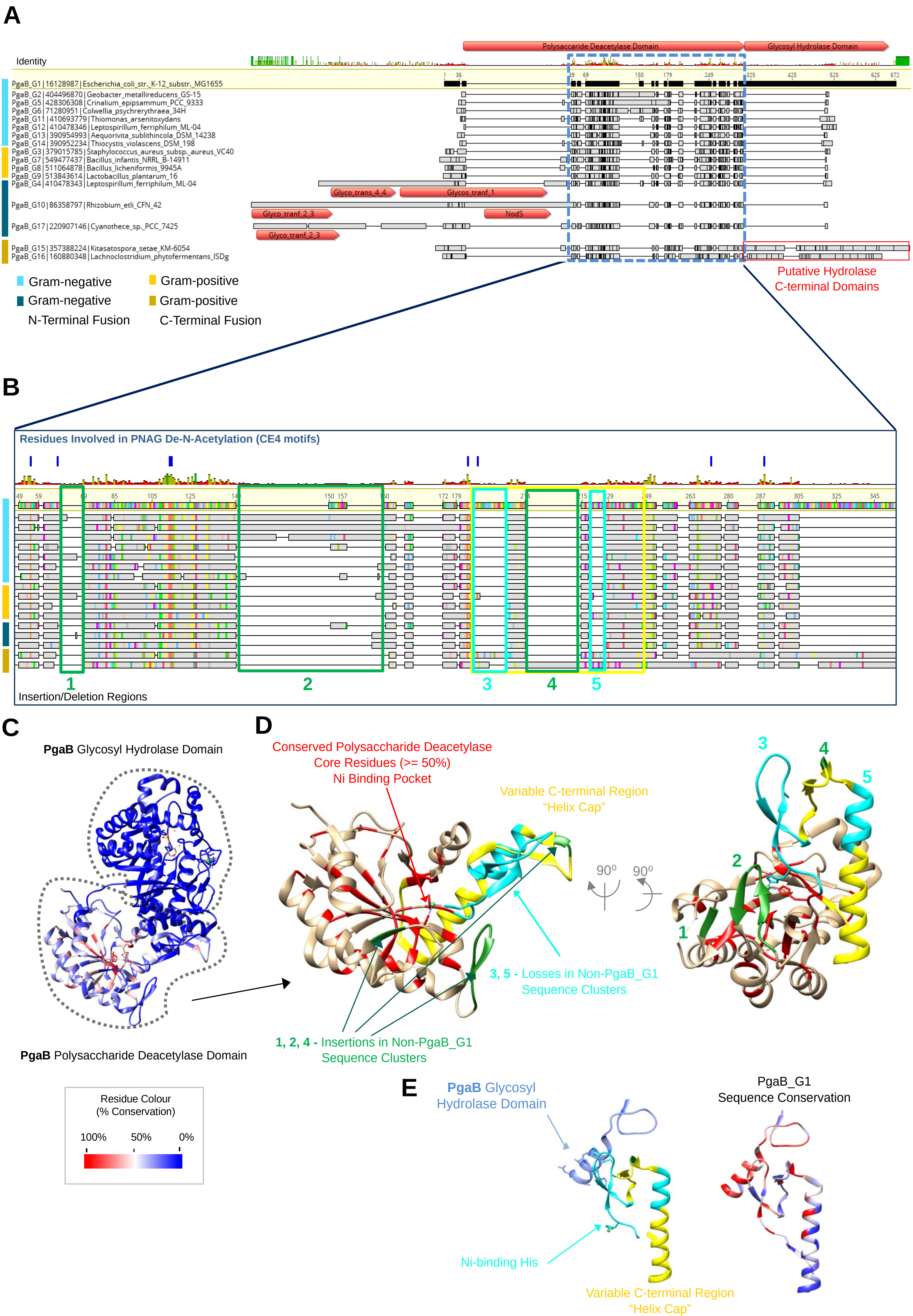
